# circPMS1-mediated LIM protein scaffolding promotes cytoskeletal remodeling and melanoma metastasis

**DOI:** 10.64898/2026.06.18.733219

**Authors:** Nicol Mecozzi, Olga Vera, Kelly M. Guzman, Manon Chadourne, Xiaonan Xu, Kaizhen Wang, Michael Martinez, Angelo A. Nicolaci, Nhan Phan, Jiqiang Yao, Margaret A. Park, Xiaoqing Yu, Alex M. Jaeger, Florian A. Karreth

## Abstract

Circular RNAs (circRNAs) have been implicated in diverse biological processes and diseases such as cancer; however, their functional roles in tumor biology remain incompletely understood. Here, we identify *circPMS1* as an aberrantly upregulated, pro-metastatic circRNA in melanoma. Overexpression and silencing studies revealed that *circPMS1* promotes migration and invasion of non-transformed melanocytes and melanoma cells, enhances metastasis in xenograft models, and drives spontaneous dissemination and lymph node metastases in a genetically engineered melanoma mouse model. *circPMS1* pro-metastatic activity depends on its circularization and is mediated by a putative secondary structure spanning the back-splice junction and the canonical start codon of the linear *PMS1* mRNA. Proteomic analysis identified LIM domain proteins, including ABLIM1, LMO7, and PDLIM7, as *circPMS1*-bound factors. Mechanistic studies demonstrate that *circPMS1* scaffolds these LIM proteins, enhancing their association with actin filaments inducing focal adhesion and cytoskeletal remodeling. This remodeling is associated with RhoA activity, and inhibition of the RhoA pathway attenuates *circPMS1*-induced migration. Together, these findings establish *circPMS1* as a pro-metastatic circRNA that drives melanoma progression by scaffolding LIM domain proteins and regulating cytoskeletal dynamics and cell motility.

## Introduction

Cutaneous melanoma is an aggressive cancer that arises from pigment-producing melanocytes and that invades surrounding tissues and metastasizes early in the course of the disease. While surgical removal of localized primary lesions is often curative, patients with melanoma that has spread to distant sites face a five-year survival rate of only ∼35%. Two key biological features are hallmarks of melanomagenesis: an exceptionally high somatic mutation burden promoting neoantigen formation^1^, and frequent hyperactivation of the MAPK signaling pathway^2,3^. These characteristics have led to the development of therapeutic strategies, including immune checkpoint blockade and MAPK targeted inhibitors. Although such treatments have significantly improved survival for individuals with advanced disease, intrinsic and acquired treatment resistance continues to limit their long-term effectiveness. Therefore, considerable research has focused on uncovering the molecular processes that drive melanoma progression and metastasis to support the development of improved therapeutic approaches. Somatic mutations in protein-coding genes commonly deregulate several critical cellular pathways, including MAPK signaling, the PI3K/AKT pathway, cell-cycle regulation, the p53 pathway, and mechanisms controlling telomerase activity^2–5^. However, these mutations occur early in tumor development, indicating that somatic mutations alone do not fully explain the transition to metastatic melanoma^3,6,7^. Recent studies have emphasized the contribution of non-mutational mechanisms to melanoma progression. These include broader genomic alterations^7^, epigenetic changes that influence gene expression^3^, and shifts in transcriptional regulation^8^. Moreover, growing evidence implicates dysregulation of non-coding RNAs in melanoma development and progression^9,10^.

Circular RNAs (circRNAs) are emerging as non-coding RNAs with critical roles in disease etiology. circRNAs exhibit a covalently closed loop configuration that lacks conventional RNA features such as 5′ and 3′ termini, a poly(A) tail, and a 5′ cap. They are generated from precursor messenger RNAs (pre-mRNAs) via a non-canonical back-splicing mechanism^11,12^ where the 3′ end of a downstream exon is spliced to the 5′ end of an upstream exon, producing a circular RNA molecule and a corresponding linear transcript lacking the circularized exon(s). Although back-splicing is thought to occur alongside canonical splicing, the determinants that favor one pathway over the other remain unclear, and the two processes may proceed in a stochastic or potentially coordinated manner^11,13^.

The circular configuration of circRNAs confers remarkable stability, as it reduces susceptibility to degradation by exonucleases and microRNA-mediated decay^11,12^. As a result, circRNAs often exhibit extended half-lives and can accumulate to substantial levels, in some cases becoming predominant transcript isoforms. Nevertheless, most circRNAs are present at lower abundance than their linear mRNA counterparts, likely reflecting the relatively lower efficiency of their biogenesis compared to canonical splicing^11,12^. Despite their generally lower abundance, circRNAs play important roles in biological processes and diseases. For example, circRNAs may function as miRNA sponges^14–18^, scaffolds for RNA binding proteins (RBPs)^19–24^, and enhancers of the expression or splicing of their cognate parental genes^25–27^, while others can encode tumor-suppressive or oncogenic peptides^28–38^.

However, despite the acknowledged functional diversity of circRNAs, their roles and mechanisms of action in melanoma remain poorly characterized. In this study, we identified *circPMS1* as a pro-metastatic circRNA that directly binds and scaffolds the LIM-domain proteins ABLIM1, LMO7, and PDLIM7 to actin filaments to promote cytoskeleton remodeling, cell motility, and melanoma metastasis formation.

## Materials and Methods

### Cell lines

Human melanocyte cell lines (Hermes1, Hermes2, Hermes3A, and Hermes4B) were obtained from St. George’s University of London (https://www.sgul.ac.uk/about/our-institutes/neuroscience-and-cell-biology-research-institute/genomics-cell-bank/cell-bank-holdings#Humanimmortalmelanocytes). Melanocytes were cultured in RPMI 1640 medium (Lonza) supplemented with 10% fetal bovine serum (FBS), 200 nM 12-O-tetradecanoylphorbol-13-acetate (TPA), 10 nM endothelin-1, 200 pM cholera toxin, and 10 ng/mL human stem cell factor. Hermes1 and Hermes3A cells expressing BRAF^V600E^ were previously generated in our laboratory^39^ and maintained in melanocyte medium lacking TPA. Melanoma cell lines, including A375 and SKMel28, were purchased from ATCC (Manassas, VA, USA). Additional melanoma lines (WM164, WM35, WM793, 1205Lu, WM115, WM266.4, and 451Lu) were generously provided by Dr. Meenhard Herlyn (Wistar Institute, Philadelphia, PA, USA) as part of the Wistar Collection of Melanoma Cell Lines. SbCl2 cells were obtained from Dr. David Tuveson (Cold Spring Harbor Laboratory, Cold Spring Harbor, NY, USA). Mouse melanoma cell lines NCC (M133M1), NPP (M171M1), BPP (M10M1), BCC (M167M1) and BCC (M161M1) were previously generated in our laboratory^40^ and cultured in RPMI 1640 supplemented with 5% FBS at 37°C with 5% CO_2_. HEK293T Lenti-X cells (Takara) were grown in DMEM (VWR) supplemented with 10% FBS. Melanocytes were maintained at 37°C in a humidified incubator with 10% CO_2_, whereas melanoma and HEK293T cells were cultured at 37°C with 5% CO_2_. All cell lines were routinely screened for mycoplasma contamination using the MycoAlert Mycoplasma Detection Kit (Lonza), and human melanoma cell lines were authenticated by short tandem repeat (STR) profiling at Moffitt’s Molecular Genomics Core.

### Plasmids, viral constructs and siRNA transfection

The ATP1-spGFP and ATP1-circPMS1 transposons were generated as described in N. Mecozzi et al., 2022^41^. The pLP-Cbh-ABLIM1-Halo, pLP-Cbh-LMO7-Halo, and pLP-Cbh-PDLIM7-Halo constructs were synthetized by Vector Builder. Mutation of the Splice Acceptor (ΔSA) or Splice Donor (ΔSD) in the ATP1-circPMS1 construct and mutation of the ATG start codon to GCG (ΔATG) in the ATP1-circPMS1 construct were done by Site Directed Mutagenesis (SDM) (New England Biolabs, Cat# E0554S) but using 2X Platinum SuperFi II Green PCR Master Mix (Invitrogen, Cat# 12369010). To test *circPMS1* IRES activity, the predicted circPMS1-IRES sequence, its antisense, and a known EMCV IRES sequence were cloned into psiCheck2 plasmid via InFusion cloning (Cat# 639650). For CasRX mediated silencing, guides targeting *circPMS1* back-splice junction were designed according to the protocol published by Zhang Y. et al., *Genome Biology* 2021^42^, and Konermann S., et al., *Cell* 2018^43^. U6-Cas13d from Addgene plasmid #166867 was cloned into a pLenti-GFP-blasticidin replacing GFP with U6-Cas13d by InFusion cloning. The guides were cloned by SDM in pLenti-U6-Cas13d-Blast lentiviral construct downstream of the Cas13d cDNA. The guide sequences are reported in **Supplementary Table 1**. For the MS2 pulldown, the MS2 motif concatemer sequence was amplified by PCR from pCDNA-MS2 tag and cloned by InFusion into the ATP1-circPMS1 transposon between circPMS1 exon 3 and 4. Furthermore, pCDNA-MS2bp-GST was obtained by cloning GST tag from pGEX-6P-1 plasmid by InFusion into pCDNA-MS2bp to replace the YFP tag. Cod_circ-v5 was cloned by inserting the linearized *circPMS1* putative coding sequence into a p-Lenti-puro construct by InFusion and the 3’ terminal V5 tag was added by SDM. For virus production, HEK239T cells were transfected with the lentiviral vector along with packaging and helper plasmids Δ8.2 and VSVG at a ratio 9:8:1 using jetPRIME transfection reagent (Polypus, Cat# 101000046). Lentiviral supernatants were collected 48 hours after transfection followed by centrifugation and filtration through a 0.45 μm syringe filter. For siRNA transfections, cells were transfected with 25 nM of either non-targeting siRNA (SMARTPool, Horizon Discovery Cat# D-001810-10-05) or sicircPMS1 (**Supplementary Table 1**), siABLIM1 (SMARTPool, Horizon Discovery Cat# L-011320-00-0005), siLMO7 (SMARTPool, Horizon Discovery Cat# L-019252-00-0005), or siPDLIM7 (SMARTPool, Horizon Discovery Cat# L-013081-00-0005) using jetPRIME transfection reagent overnight. The medium was refreshed after 24 hours, and the cells were collected 72 hours after transfection for protein and RNA isolation.

### Treatments

*RNase R:* 2 μg of RNA sample were treated with 20 U/μL of RNaseR (Biosearch Technologies, E0111-20D1, SS000769-D1) following manufacturer’s instructions at 37°C for 30 minutes. RNA was isolated again after treatment.

*Actinomycin D:* 5 μg/μL of Actinomycin D (Cell Signaling, Cat# 15021S) was used to treat 500,000 melanoma cells for 4-8-12-16 hours.

*Rhosin:* 30 μM of Rhosin (TOCRIS, Cat# 1281870-42-5) were used to treat melanoma cells for 96 hours. DMSO was used as control.

*CN03:* 1μg/mL of CN03 (Cytoskeleton, Cat# CN03-A) was used to treat melanoma cells for 4 hours. DMSO was used as control.

*RNase 1/A:* Cell lysates were treated with 100 μg/mL of RNase A (Thermo Scientific Cat# EN0531) and 20 U of RNase I (Thermo Fisher Scientific, Cat# EN0601) per sample at 37°C for 30 minutes.

### Proliferation assay

1,000-2,500 cells were seeded in 96-well plates in triplicates and cultured for 3-5 days. Cell proliferation was measured using the CellCyte X instrument (Cytena), analyzing the total cell confluency over time.

### Focus formation assay

A total of 700-2,000 cells per well were seeded in 6-well plates and maintained in culture for 14-25 days. Colonies were fixed with methanol and stained for 20 minutes using 0.1% crystal violet (VWR, Cat# 97061–850) prepared in 20% methanol. Following staining, an equal volume of 10% acetic acid was added to each well to solubilize the dye. Colony formation was quantified by measuring the absorbance of the resulting solution at 600 nm.

### Transwell assay

For migration assays, 10,000-30,000 cells in 2.5% FBS media were seeded in transwells and cultured for 24 hours in wells containing 15% FBS media. For invasion assays, the same procedure was followed but transwells were coated with 25 μL of Matrigel (VitroGel, Cat# VHM01) diluted 1:4 in DMEM media and the cells were cultured for 48 hours. At endpoint, transwell membranes were fixed with Paraformaldehyde solution, 4% in PBS (Thermo Fisher Scientific, Cat# J19943-K2) for 10 minutes and stained with a 0.1% crystal violet (VWR, Cat# 97061–850) solution prepared in 20% methanol for 20 minutes. The bottom of the transwell membranes were imaged using Zeiss Axiocam 305 color. Cells were counted using on ImageJ in 3-5 fields per membrane.

### Cell Death assay

Cell death was assessed using SYTOX Green dye (Thermo Fisher, Cat# S7020). SYTOX Green (20 nM) was added to the culture medium, and cell death was monitored over 3 days. Cell viability and death were quantified using a Cellcyte X instrument (Cytena) by normalizing the number of green fluorescent objects (dead cells) to cell confluency. Data is plotted at 36 hours after treatment.

### Animal studies

All animal studies were performed in accordance with a protocol approved by the Institutional Animal Care and Use Committee at the University of South Florida (Tampa, FL). NSG mice (stock no. 005557) were purchased from The Jackson Laboratory (JAX). For xenograft studies, mice were anesthetized with isoflurane, and 1×10^5^ luciferase-positive cells were administered via tail vein injection. Metastatic burden was monitored weekly using an IVIS imaging system (Revvity). Mice were euthanized upon reaching defined humane endpoints. Organs were subsequently collected, fixed in 10% formalin, and processed for hematoxylin and eosin staining.

Derivation and validation of embryonic stem (ES) cells on a mixed C57BL/6 x 129Sv background was performed as previously described^44^. Braf^CA^; Pten^FL/FL^; TyrCre-ERt2; CAGs-LSL-rtTA3; CHC (BPP) ES cells were targeted with the TRE-circPMS1 construct by recombination-mediated cassette exchange as described previously^44^, and experimental chimeras were generated in Moffitt’s Gene Targeting Core. Melanoma was initiated by applying 1 µl of a 25 mg/ml solution of 4-Hydroxytamoxifen (4-OHT, Millipore Sigma, Cat# H6278) in DMSO either on a single spot of the shaved backs or by dipping the tails in 25 mg/ml of 4-OHT of 3 weeks old mice. Immediately following 4-OHT application, chimeras were either placed on a 200 mg/kg doxycycline diet (Envigo) or left on a normal chow diet as control. Mice were euthanized once the tumors reached IACUC-approved endpoints. Tumors, mandibular and inguinal lymph nodes were collected for further analysis.

### Fluorescent In situ hybridization (FISH)

25,000 melanoma cells were plated in 8-chamber culture slides (Falcon, Cat# 354108). After 24 hours, FISH was performed according to published methods^45^ with the following modifications. Culture media was removed and the cells were washed twice with 1X PBS, followed by 10 minutes of fixation at room temperature (3.7% formaldehyde in 1X PBS). Cells were washed in 1X PBS followed by permeabilization (1% TritonX-100 in 1X PBS) for 30 minutes at room temperature and treatment with 12.5 U/mL of RNase R for 30 minutes at 37°C. Cells were washed in 1X PBS and incubated with 125 nM of *circPMS1* probe (**Supplementary Table 1**, Dharmacon) in 100 μL of hybridization buffer (10% w/v dextran sulfate and 10% formamide in 2X SSC) at 37°C overnight. The following day, cells were washed with wash buffer (10% formamide in 2X SSC) for 30 minutes at 37°C and with 1X PBS and then incubated with 1:100 of 400X Alexa Fluor 546 Phalloidin (Invitrogen, Cat# A22283) for 45 minutes at room temperature, followed by 3 washes in 1X PBS for 3 minutes each. Slide were mounted using ProLong Diamond Antifade mounting media containing DAPI (Invitrogen, Cat#P36971). Cells were imaged using the Leica Microsystems Confocal SP8 microscope. FISH on tissue was performed according to published methods^46^ with the following modifications. The slides were deparaffinized and dehydrated with two washes in xylene for 5 minutes each, two washes in 100% ethanol for 3 minutes each, one wash in 95% ethanol for 3 minutes, one wash in 70% ethanol for 3 minutes. Everything else was performed as per protocol until point 4 of the “Post-hybridization washes” section. The analysis was performed with ImageJ.

### Immunofluorescence (IF)

25,000 melanoma cells were plated in 8-chamber culture slides. After 24 hours, culture media was removed and cells were washed twice with 1X PBS, followed by 10 minutes of fixation at room temperature (0.1% Triton-X in 4% Paraformaldehyde in PBS solution). Slides were rinsed twice in 1X PBS for 5 minutes each. Permeabilization was performed using 1% Triton x-100 in PBS for 30 min at room temperature. Slides were rinsed again twice in 1X PBS for 5 min each. Cells were then incubated in 10% normal goat serum in 1X PBS for 30 min at room temperature. Cells were then incubated with primary antibody (**Supplementary Table 2**) at 1:50-1:100 dilution in 1X PBS overnight at 4°C. On the following day, cells were rinsed three times in 1X PBS for 5 min each and incubated with fluorochrome-conjugated secondary antibody at 1:1000 dilution in 1X PBS for 1 hour at 37°C in the dark. Cells were rinsed three times in 1X PBS for 5 min each in the dark and incubated with 1:100 Phalloidin (Invitrogen, Cat# A22283) for 45 minutes at room temperature, followed by 3 washes in 1X PBS for 3 minutes each. Slides were mounted using ProLong Diamond Antifade mounting media containing DAPI (Invitrogen, Cat#P36971). Cells were imaged using the Leica Microsystems Confocal SP8 microscope.

For IF on tissue, slides were deparaffinized and dehydrated with two washes in xylene for 5 minutes each, two washes in 100% ethanol for 3 minutes each, one wash in 95% ethanol for 3 minutes, one wash in 70% ethanol for 3 minutes, one wash in 50% ethanol for 3 minutes. The slides were incubated in 10 mM citrate buffer pH=6, boiled in the microwave for 10 minutes and cooled down at room temperature for 30 minutes. Slides were then submerged in 3% H_2_O_2_ for 10 minutes at room temperature and transferred onto tissue holder racks. Slides were washed three times with 1X PBS supplemented with 0.1% TritonX-100 (PBS-T). Tissues were blocked using 125 μL of 2.5% horse serum in PBS-T for 20 minutes at room temperature, followed by incubation with primary antibody (SOX10 1:100 dilution) at 4°C overnight. The following day, slides were washed three times with PBS-T, incubated with fluorochrome-conjugated secondary antibody at 1:1000 dilution in 1X PBS for 5 hours at room temperature in the dark. After three washes in PBS-T, slides were mounted with ProLong Diamond Antifade mounting media containing DAPI. All IF images were taken with a 63X objective. Analyses were performed with CellProfiler by identifying the nucleus, producing cell segmentation and thresholding at 15% colocalization mask identifying colocalization metrics.

### MS2 RNA pulldown

Melanoma cells were grown to confluence in 100 or 150 mm plates (Sarstedt, Cat# 83.3902; 83.3903.300), washed with 1X PBS, and UV-crosslinked at 150 mJ/cm^2^ for 30 seconds using a UVP Crosslinker (Analytik Jena). Cells were washed again, scraped, and collected into tubes containing 100 μL lysis buffer (50 mM Tris, pH 7.5; 150 mM NaCl; 1 mM EDTA; 0.5% NP-40). Cells were lysed on ice for 20 minutes and sonicated for 1 minute. Samples were centrifuged at 15,000 rpm for 15 minutes at 4°C, and the supernatant was collected. Protein concentration was determined using a BSA assay, and 1 mg of total protein was used for downstream applications. Forty microliters of GSH-Sepharose slurry (GoldBio, Cat# G-250-5) were added to each lysate and incubated overnight at 4°C with rotation. The following day, resin–lysate mixtures were centrifuged at 5,000 rpm for 1 minute at 4°C, and the supernatant was discarded. The resin was washed once with wash buffer 1 (50 mM Tris, pH 7.5; 150 mM NaCl; 1 mM EDTA; 0.05% NP-40), followed by centrifugation at 5,000 rpm for 1 minute at 4°C. This wash step was repeated five additional times. A final wash was performed using wash buffer 2 (50 mM Tris, pH 7.5; 150 mM NaCl; 1 mM EDTA). After removing the final wash, 70 μL of elution buffer (50 mM Tris, pH 7.5; 150 mM NaCl; 1 mM EDTA; 20 mM reduced glutathione) was added, and samples were incubated for 20 minutes at room temperature. The resin was centrifuged at 5,000 rpm for 1 minute, and the supernatant was collected. For RNA isolation, the eluate was treated with wash buffer 2 supplemented with proteinase K and incubated for 20 minutes at 55°C followed by 10 minutes at 95°C. For protein analysis, the eluate was mixed with sample buffer (20% glycerol, 6% SDS, 0.125 M Tris, pH 6.8) and heated at 95°C for 10 minutes. The conditions used for MS2 pulldown and Mass Spec sequencing were MS2 only, *circPMS1*-MS2 and *circPMS1* only.

### Halo-tag pulldowns

Melanoma cells were grown to confluence in 100 or 150 mm plates, washed with 1X PBS, and UV-crosslinked at 150 mJ/cm^2^ for 30 seconds (no crosslinking was performed for Halo-tag pulldowns upon Rhosin or CN03 treatment). Cells were washed again, scraped, collected into tubes, and centrifuged at 13,000 rpm for 5 minutes. The PBS was replaced with 1X RIP buffer (50 mM Tris-HCl, pH 7.5; 10 mM EDTA; 1.5 M NaCl; 5% IGEPAL), and samples were incubated on ice for 30 minutes with vortexing every 5 minutes. Lysates were then centrifuged at 13,000 rpm for 10 minutes, and the supernatant was collected. Meanwhile, 50 μL HaloTag magnetic beads (Promega, Cat# G7281) were washed with 1X PBS and blocked with 1% BSA for 20 minutes at room temperature on a rotator. Beads were subsequently washed with 1X RIP buffer. The prepared lysates were then added to the beads and incubated overnight at 4°C with rotation. On the following day, beads were washed twice for 5 minutes at 4°C with High Salt Buffer (50 mM Tris-HCl, pH 7.5; 1 mM EDTA; 1% NP-40; 0.1% SDS; 0.5% sodium deoxycholate), followed by two 10-minute washes at 4°C with IP Wash Buffer (20 mM Tris-HCl, pH 7.5; 10 mM MgCl₂; 0.4% Tween-20). For RNA isolation, beads were incubated with IP Wash Buffer supplemented with 8 U of Proteinase K (New England Biolabs, Cat# P8107S) at 50°C for 20 minutes. After incubation, beads were resuspended in TRIzol, and RNA was extracted according to standard procedures. For protein isolation, beads were incubated with 1X TEV reaction master mix (5μL of 20X ProTEV Buffer, 1 μL of 100 mM DTT, 3μL of ProTEV enzyme and water up to 100 μL) for 75 minutes at 30°C shaking at 1,200 rpm in a ThermoMixer. The reaction was stopped by adding 1X LDS sample buffer (NuPAGE, Cat# NP0008) and heated at 70°C for 10 minutes. The supernatant was collected and resolved on 4-12% NuPAGE gels (Invitrogen, Cat# WG1402B0X) for Western blot analysis.

### Active Rho pulldown

1 mg of protein from parental WM35 cells or cells overexpressing circPMS1 or spGFP was collected for pulldown with Active Rho Pull-down and detection kit (Thermo Fisher Scientific, Cat# 16116). 1 mg of WM35 was divided in half for GDP negative control and GTPγS positive control spike-in. The pulldown was performed following the manufacturer’s instructions.

### CLAP

WM35 melanoma cells stably expressing pLP-Cbh-ABLIM1-Halo, pLP-Cbh-LMO7-Halo, or pLP-Cbh-PDLIM7-Halo constructs were cultured to confluence, washed with 1X PBS, and UV-crosslinked at 150 mJ/cm^2^. Cells were scraped, collected, and centrifuged at 12,000 rpm for 5 minutes at 4°C. Cell pellets were lysed in 200 μL of RIPA buffer supplemented with 50X Promega protease inhibitor cocktail, TURBO DNase, 100X Mn^2^⁺/Ca^2^⁺ mix, and RiboLock RNase inhibitor. Lysates were incubated on ice for 20 minutes with vortexing every 5 minutes, then warmed to 37°C to activate DNase. Samples were centrifuged at 15,000 g for 20 minutes, and the supernatant was collected and subsequently treated with RNase R as described below. In parallel, 20 μL of HaloTag magnetic beads (Promega, Cat# G7281) per sample were magnetically separated and washed with 200 μL of 1X PBS containing 0.1% Triton X-100. Beads were blocked with 400 μL of 1% BSA in PBS for 20 minutes at room temperature with rotation, washed three times with 1X PBS + 0.1% Triton X-100, and kept on ice until use. Lysates were incubated with prepared beads overnight at 4°C with end-over-end rotation. The following day, beads were subjected to sequential high-stringency washes at 95°C for 3 minutes each using the following buffers: NLS CLAP Wash Buffer (600 μL of 20% NLS, 120 μL of 500 mM EDTA, 600 μL of 10X PBS and water up to 6 mL), 1 M NaCl CLAP Wash Buffer (300 μL of 1 M HEPES pH 7.4, 1200 μL of 5 M NaCl, 60 μL of 10% NP-40, water up to 6 mL), 8 M Urea CLAP Wash Buffer (300 μL of 1 M HEPES pH 7.4, 60 μL of 10% NP-40, 5640 μL of 8 M Urea), Tween-20 CLAP Wash Buffer (300 μL of 1 M HEPES pH 7.4, 60 μL of 10% NP-40, 60 μL of 10% Tween-20, water up to 6 mL), and TEV CLAP Wash Buffer (300 μL of 1 M HEPES pH 7.4, 60 μL of 10% NP-40, 12 μL of 500 mM EDTA, water up to 6 mL). For RNA isolation, beads were incubated with 100 μL of NLS CLAP Wash Buffer supplemented with 8 U of Proteinase K (New England Biolabs, Cat# P8107S) for 20 minutes at 50°C. The supernatant was collected, and RNA was extracted using TRIzol reagent. The RNA was reverse transcribed to cDNA and the qRT-PCR product was run on a 1% agarose gel.

### Proximity Ligation Assay (PLA)

PLA was performed using Duolink In situ Fluorescence (Sigma Cat# DU092004, DU092002, DU092007) according to the manufacturer’s recommendation with fixation and permeabilization of cells performed as described for IF. The primary antibody dilution was 1:50 and the pairing used were LMO7 (mouse)-PDLIM7 (rabbit), LMO7 (mouse)-ABLIM1 (rabbit), PDLIM7 (rabbit)-ABLIM1 (mouse). Analysis was performed with CellProfiler. All confocal microscope images were taken with a 63X objective.

### Western Blot

Protein samples were homogenized in RIPA buffer (50 mM Tris-HCl, pH 8.0, 1 mM EDTA, 0.5 mM EGTA, 1% Triton X-100, 0.5% sodium deoxycholate, 0.1% SDS, 150 mM NaCl) supplemented with protease and phosphatase inhibitors. For each sample, 20 μg of total protein was combined with Laemmli buffer, boiled, and resolved on NuPAGE 4-12% precast gels. Proteins were transferred onto nitrocellulose membranes using standard wet-transfer methods. Membranes were blocked for 1 hour at room temperature in 5% non-fat dry milk prepared in TBST (20 mM Tris, 150 mM NaCl, 0.1% Tween-20), followed by overnight incubation at 4°C with primary antibodies diluted in 5% milk or BSA. All antibodies used are listed in **Supplementary Table 2**. Membranes were washed three times for 5 minutes each in TBST and then incubated with horseradish peroxidase-conjugated secondary antibodies (1:10,000 dilution) for 1 hour at room temperature. After three additional washes in TBST, blots were developed using chemiluminescent substrate (1:1 dilution) for 3 minutes. Signals were detected using a ChemiDoc MP (BioRad), ProSignal Blotting Film (Prometheus, Cat# 30-810L) or LI-COR imaging system.

### RT-qPCR

Total RNA was extracted using TRIzol™ Reagent (Invitrogen, Cat#15596018) according to the manufacturer’s instructions. For each sample, 1,000 ng of RNA was reverse transcribed into cDNA using PrimeScript RT Master Mix (TakaraBio, Cat# RR036A). cDNA was diluted 1:20 and subjected to quantitative PCR using 2X Universal SYBR Green Fast qPCR Mix (ABclonal, Cat# RK21203). Reactions were performed in duplicate or triplicate on a StepOne Plus Real-Time PCR System (Applied Biosystems, Foster City, CA, USA). Relative gene expression was determined using the comparative threshold cycle (2^-ΔΔCt) method, with Actin used as the internal reference for normalization. Primer sequences used for RT-qPCR are provided in **Supplementary Table 1**.

### Luciferase assay

20,000 cells/well in six replicates per condition (circPMS1-IRES, EMCV-IRES, As_IRES) were plated in a 96-well plate and luciferase assay was performed using the Dual-Luciferase Reporter Assay System (Promega, Cat# E195A) and analyzed using the GlowMax Discover instrument (Promega).

### Mass Spectrometry

Digestion was performed with 2 mM TCEP reduction and 20 mM IAA alkylation followed by digestion with 200 ng of trypsin overnight at 37 °C. Another aliquot of 200 ng trypsin was added the next day for an additional 2-hour digest. Peptides in solution were acidified to 1% trifluoroacetic acid, then desalted using a SOLAu plate (Thermo #60209-001). Eluted, desalted peptides were assayed via a peptide concentration assay (Pierce #23275). 500 ng of peptide were vacuum-centrifuged to dryness and resuspended in 30 mL of 0.1% trifluoroacetic acid, then loaded on EvoSep EvoTip Pure tips in preparation for LC-MS/MS analysis. A nanoflow ultra high-performance liquid chromatograph and nanoelectrospray orbitrap mass spectrometer (EvoSep and Q-Exactive plus) were used for LC-MS/MS. The sample was loaded onto an EvoTip pure (EV2013). Trapped peptides were eluted onto the analytical column (EV1106, 15 cm length x 150 µm ID, 1.9 µm particle size). A factory default extended gradient (88-minute) was used with solvent A (water + 0.1% formic acid) and solvent B (acetonitrile + 0.1% formic acid). Spray voltage was 1900 V. Capillary temperature was 275°C. S lens RF level was set at 50. Data-dependent acquisition was performed using Top16 precursors. The resolution for MS and MS/MS were set at 70,000 and 17,500 respectively. Dynamic exclusion was 15 seconds for previously sampled peaks.

DDA MS searching was performed with FragPipe (v.21.0) using the Label Free Quantification with Match Between Runs workflow against the Gencode proteome (v42) with group-specific FDR set to 1%. The search was completed with tryptic digestion with allowed missed cleavages set to 2. Methionine oxidation, cysteine carbamidomethylation and protein N-term acetylation were included as variable modifications, and peptide length 7-50 amino acids. Label free quantification was performed using IonQuant using precursor abundance. Match between runs (MBR) was set to true with an FDR of 1%. Search outputs were subsequently imported into R for further analysis. Proteins that exhibited >1.5 fold enrichment in circPMS1-MS2 pulldowns versus MS2 only or circPMS1 only controls in both A375 cells and WM35 cells were prioritized for further validation.

### RNA sequencing

circRNA expression analysis was performed on previously published paired-end RNA-seq data from melanoma and melanocyte cell lines (GSE148552). Sequencing adapters and low-quality bases were removed by adapter and quality trimming prior to alignment. Reads were then aligned to the human reference genome (hs37d5) using STAR^47^ with chimeric read detection enabled (--chimSegmentMin 10). circRNAs were identified and annotated by CIRCexplorer2 using GENCODE v25lift37 annotation based on back-splice junction reads. circRNA junction counts from all samples were merged into a count matrix. Differential expression analysis between melanoma and melanocyte cell lines was performed using DESeq2, and significance was determined using Benjamini-Hochberg adjusted p-values^47–49^. To assess the ratio of circular to linear RNA, the DCC/CircTest pipeline was used. CircRNAs were detected using the DCC tool with the requirement that at least 5 reads were detected in at least 6 samples. Circular to linear ratios and global p-values for the groups were calculated using CircTest^50^. Beta-binomial model with a Benjamini-Hochberg adjustment was used to test for differences in melanoma versus melanocyte cell lines. Because CIRCexplorer2 (GRCh37/hg19) and the DCC/CircTest pipeline (GRCh38/hg38) report back-splice junctions in different genome assemblies, DCC/CircTest coordinates were converted from GRCh38 to GRCh37 using the UCSC liftOver tool. Lifted coordinates were matched to CIRCexplorer2-derived circRNAs by genomic position, allowing a tolerance of up to 2 bp at each junction boundary to accommodate differences in back-splice junction coordinate conventions between the two tools.

For mRNA expression analysis by RNA-seq, WM35 melanoma cells were cultured in 10 cm plates and RNA was isolated in triplicates using the miRNeasy Kit (Qiagen, Cat# 217004). RNA was quantitated with the Qubit Fluorometer (ThermoFisher Scientific, Waltham, MA) and screened for quality on the Agilent TapeStation 4200 (Agilent Technologies, Santa Clara, CA). The samples were then processed for RNA-sequencing using the NuGEN Universal RNA-Seq Library Preparation Kit (Tecan Genomics, Redwood City, CA). Briefly, 100 ng of RNA was used to generate cDNA and a strand-specific library following the manufacturer’s protocol. Quality control steps were performed, including TapeStation size assessment and quantification using the Kapa Library Quantification Kit (Roche, Wilmington, MA). The final libraries were normalized, denatured, and sequenced on the Illumina NovaSeq 6000 sequencer with the SP-200 cycle reagent kit in order to generate approximately 80 million 101-base read pairs per sample (Illumina, Inc., San Diego, CA).

Raw sequencing reads were processed for adapter detection and removal using BBMerge (v37.02) and cutadapt (v1.8.1), respectively. Processed reads were mapped to the GRCh38 human reference genome using STAR (v2.7.7a). Gene-level expression quantification was performed with RSEM (v1.3.0) based on GENCODE v30 annotations. Statistical normalization and differential expression analyses across experimental groups were conducted using DESeq2^47,49,51–53^.

### Statistical analysis

Results are expressed as mean ± standard error of the mean (SEM). All experiments were conducted 2-5 independent biological replicates, each including 2-3 technical replicates; one representative experiment is presented. Statistical differences between two groups were evaluated using an unpaired, two-tailed t-test. A p-value < 0.05 was considered statistically significant.

### Data availability

Sequencing data supporting the findings of this study have been deposited in the Gene Expression Omnibus under accession numbers GSE335105 (reviewer token: wvchsqaetravbch) (RNA-seq of circPMS1 overexpression in WM35 cells). Mass spectrometry proteomics data have been deposited to the ProteomeXchange Consortium via PRIDE and can be accessed through accession number PXD079605 (reviewer token: K5pcsAE3spXG). Previously published dataset GSE148552 (RNA-seq of melanocyte and melanoma cell lines) is available at GEO under https://www.ncbi.nlm.nih.gov/geo/query/acc.cgi?acc=GSE148552.

## Results

### Identification of differentially expressed circRNAs in melanoma cells

To identify differentially expressed circRNAs in melanoma, we analyzed an RNA-sequencing dataset (GSE148552) consisting of four human melanocyte cell lines (Hermes1, Hermes2, Hermes3A, and Hermes4B) and five human melanoma cell lines (1205Lu, SKMel28, WM164, WM35, and WM793) using CIRCexplorer2 and DESeq2. This analysis identified a total of 6,459 circRNAs, of which 86 were significantly differentially expressed (padj < 0.05 and |log2FC| >2) in melanoma relative to melanocytes (**Fig. 1A, Supplementary Table S3**). circRNA levels may change due to altered transcription or splicing regulation. Changes in transcription affect the parental mRNA and its encoded protein, and an accompanying change in circRNA abundance may therefore be a by-product with no impact on melanoma biology. In contrast, changes in back-splicing can alter circRNA levels with relatively little effect on the parental mRNA, given its typically greater abundance. We therefore sought to distinguish circRNAs whose abundance change is accompanied by a shift in circularization from those that simply track host-gene transcription. To this end, we analyzed the same dataset with the DCC/CircTest pipeline, which quantifies the ratio of circular to cognate linear junction reads. This detected 359 circRNAs, 23 of which were among the 86 differentially abundant circRNAs (**Supplementary Table S4**). For these 23 circRNAs, we compared their change in abundance (DESeq2 log2 fold change) with their change in circularization (CircTest log2 odds ratio of the circular-to-linear ratio; **Fig. 1B**). This comparison was restricted to the differentially abundant circRNAs detected by CircTest (22 of 23 with an estimable circularization odds ratio); for the remaining 63, circularization could not be assessed, and whether their abundance changes reflect altered back-splicing or transcription is unknown.

**Fig. 1.**
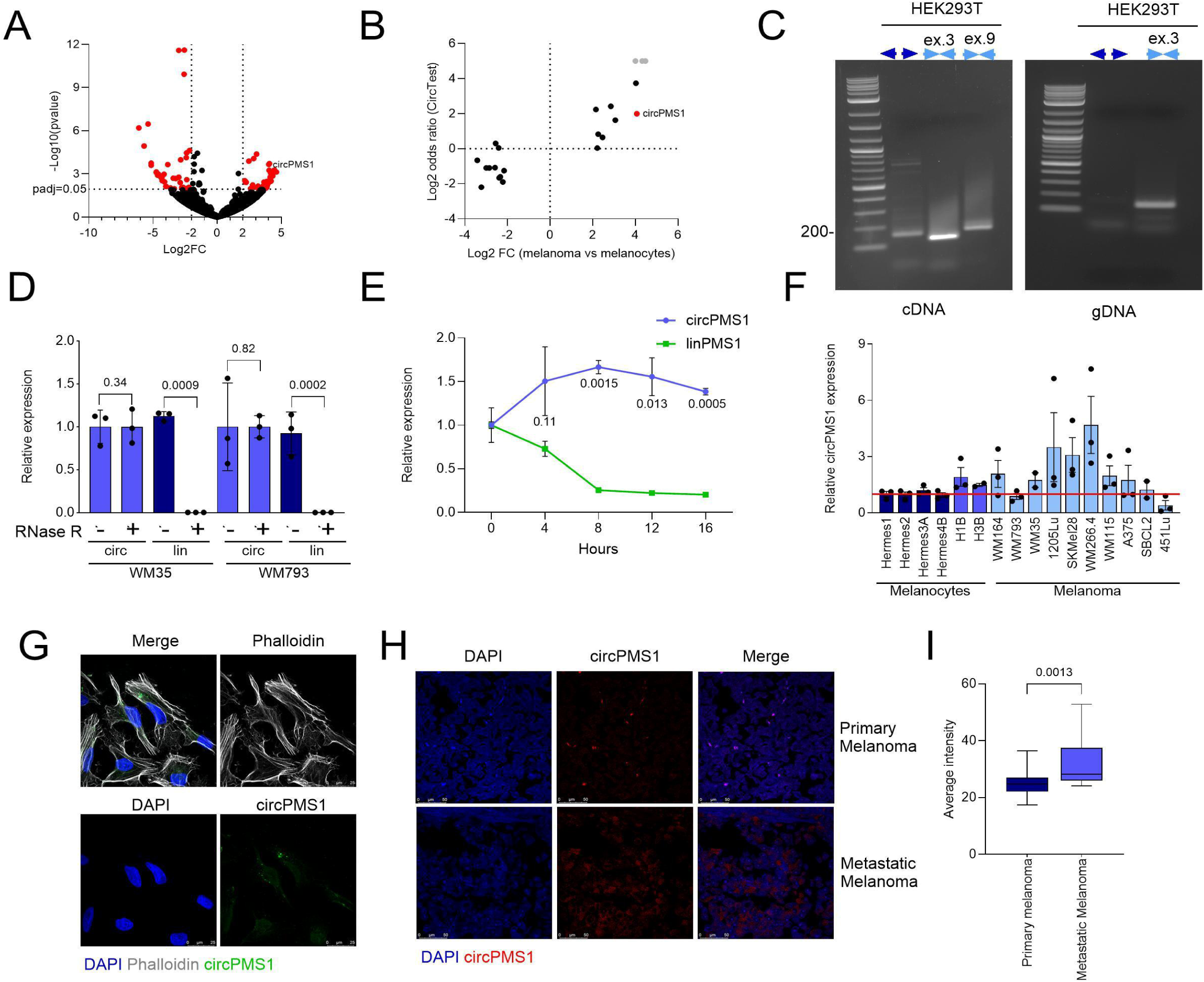
Identification and characterization of *circPMS1*. (**A**) Volcano plot representing all identified circRNAs by RNA-seq. |Log_2_FC| >2 and padj<0.05 were used as selection criteria. 86 circRNAs are significantly differentially expressed. (**B**) Comparison of abundance and circularization changes for the 23 differentially abundant circRNAs detected by both CIRCexplorer2/DESeq2 and the DCC/CircTest pipeline. Each point represents one circRNA, positive values indicate higher abundance or circularization in melanoma. *circPMS1* is shown in red. Gray symbols denote circRNAs with zero circular fraction in melanocytes, for which the odds ratio is undefined and capped; LPAR1 was undetectable in melanocytes and is not shown. No significance threshold is applied; circularization differences were modest and none reached statistical significance after correction for multiple testing, likely reflecting the limited number of cell lines and the variability in circularization between them (see Supplementary Table S4). (**C**) Left: Agarose gel showing RT-qPCR products using cDNA from HEK293T cells as template. *circPMS1* was amplified using divergent primers across the back-splice junction or convergent primers in exon 3 and *PMS1* was amplified with convergent primers in exon 9. Right: Agarose gel showing PCR products using genomic DNA from HEK293T cells as template and the *circPMS1* divergent or exon 3 convergent primers. (**D**) RT-qPCR showing *circPMS1* and linear *PMS1* expression in WM35 and WM793 cells. Lysates were treated with RNase R or untreated (**E**) RT-qPCR showing *circPMS1* and linear *PMS1* half-life upon Actinomycin D treatment. (**F**) RT-qPCR of endogenous *circPMS1* expression in a panel of melanocytes (dark blue), melanocytes harboring BRAF^V600E^ (medium blue), and melanoma cell lines (light blue). *circPMS1* expression was normalized to Hermes1 (depicted by the red line). (**G**) Fluorescent in situ hybridization for *circPMS1* in 1205Lu cells. (**H**) Quantification of *circPMS1* fluorescent in situ hybridization signal in a melanoma tissue microarray (n= 5 fields). Statistical significance was determined using t-test.

### *circPMS1* emerged as the most compelling candidate

It exhibited among the greatest increases in abundance (log2FC ∼4), comparable to *circCDK13*; (**Supplementary Table S3**); however, the *CDK13* locus produces at least two distinct circular isoforms (**Supplementary Table S3**), whereas the *PMS1* locus generates a single circRNA consistently expressed in melanoma cells, making *circPMS1* a more tractable candidate for functional study. *circPMS1* also showed a trend toward an increased circular fraction in melanoma. On the basis of its strong upregulation and its origin as a single dominant isoform, we selected *circPMS1* for in-depth investigation.

*circPMS1* is a 602 nucleotide circRNA, generated from exons 2-5 of the *PMS1* gene located on chromosome 2. In melanoma cells, 5-13% of the *PMS1* transcripts are circularized (**Supplementary Fig. S1A**). To validate *circPMS1* expression and discriminate it from the linear *PMS1* transcript, divergent primers were designed such that the forward primer annealed to exon 5 and the reverse primer to exon 2 of *circPMS1* (**Supplementary Fig. S1B**). Using cDNA from HEK293T cells as template, these divergent primers selectively amplified *circPMS1*, as confirmed by the expected amplicon size, whereas *PMS1* was amplified only by convergent primers targeting either exon 3, shared with *circPMS1*, or exon 9, which is exclusive to the linear transcript (**Fig. 1C**). Importantly, divergent primers failed to amplify a product when HEK293T genomic DNA was used as a template, whereas convergent exon 3 primers successfully amplified the genomic *PMS1* sequences (**Fig. 1C**). The specificity of the divergent primers was further validated by Sanger sequencing of the back-splice junction (**Supplementary Fig. S1C**), which also confirmed the expected linkage of exon 5 with exon 2. Given their circular nature, circRNAs are resistant to exonuclease-mediated degradation. *circPMS1* levels were stable when RNA from WM35 and WM793 melanoma cells was treated with RNase R, while *PMS1* levels were markedly reduced (**Fig 1D**). Moreover, treatment of WM35 cells with actinomycin D demonstrated significantly greater transcript stability for *circPMS1* than *PMS1* (**Fig. 1E**), supporting the notion that *circPMS1* is a circular RNA.

Expression analysis by RT-qPCR across a panel of melanocyte and melanoma cell lines validated significant upregulation of *circPMS1* in melanoma cells (**Fig. 1F**). *PMS1* showed more variable expression, with increased expression in some melanoma cell lines compared to melanocytes (**Supplementary Fig. S1D**). *circPMS1* expression was also detected in additional cancer cell lines, including pancreatic (T3M4 and PATU), ovarian (OVC5 and OVC8), and lung cancer (A549) (**Supplementary Fig. S1E**). Divergent primers for the murine analog detected *circPMS1* expression also in mouse embryonic fibroblasts, with a back-splice junction between exons 2 and 5 (**Supplementary Fig. S1F, G**). *circPMS1* expression was also detected across multiple murine tissues and organs, with the highest levels found in the brain and kidneys (**Supplementary Fig. S1H**), indicating that *circPMS1* is conserved in mice.

Subcellular localization analyses performed by fluorescent in situ hybridization (FISH) revealed that endogenous *circPMS1* was present in both the cytoplasmic and nuclear compartments of melanoma cells (**Fig. 1G**). Notably, FISH analysis of melanoma tissue microarrays (TMAs) demonstrated significantly higher *circPMS1* expression in metastatic melanoma samples relative to primary melanomas (**Fig. 1H, I**), suggesting a potential role for *circPMS1* in melanoma progression and metastasis. Collectively, these results identify *circPMS1* as a broadly expressed and conserved circRNA that is aberrantly upregulated in melanoma.

### *circPMS1* promotes cell migration and invasion *in vitro*

To investigate whether *circPMS1* has a pro-tumorigenic role in melanoma, we delivered ectopic *circPMS1* or a circular GFP control (spGFP) using a transposon approach that enables stable overexpression of circRNAs^41^. Ectopic *circPMS1* exhibited a subcellular localization pattern similar to endogenous *circPMS1* (**Supplementary Fig. 1I**). Overexpression of *circPMS1* in human melanoma cell lines exhibiting low-to-moderate endogenous *circPMS1* expression (WM35, WM164, WM793, A375) (**Supplementary Fig. S2A**) had no effect on proliferation or low-density focus formation (**Supplementary Fig. S2B-G**). In contrast, *circPMS1* overexpression markedly enhanced migration (**Fig. 2A**) and invasion (**Fig. 2B**) of these melanoma cells in transwell assays. The effects of *circPMS1* are not restricted to malignant cells as ectopic expression of *circPMS1* in wildtype (Hermes1) and BRAF^V600E^-mutant (H3B) human melanocytes increased migration and invasion (**Fig. 2C-F, Supplementary Fig. S2H, I**). Moreover, ectopic *circPMS1* expression in four murine melanoma cell lines with different mutational backgrounds (BRAF^V600E^; PTEN^-/-^ (BPP), BRAF^V600E^; CDKN2A^-/-^ (BCC), NRAS^Q61R^; PTEN^-/-^ (NPP), or NRAS^Q61R^; CDKN2A^-/-^ (NCC)) (**Supplementary Fig. S2J**) also enhanced migration and invasion (**Supplementary Fig. S2K, L**), suggesting that the effect of *circPMS1* is conserved across species and independent of driver mutations status.

**Fig. 2.**
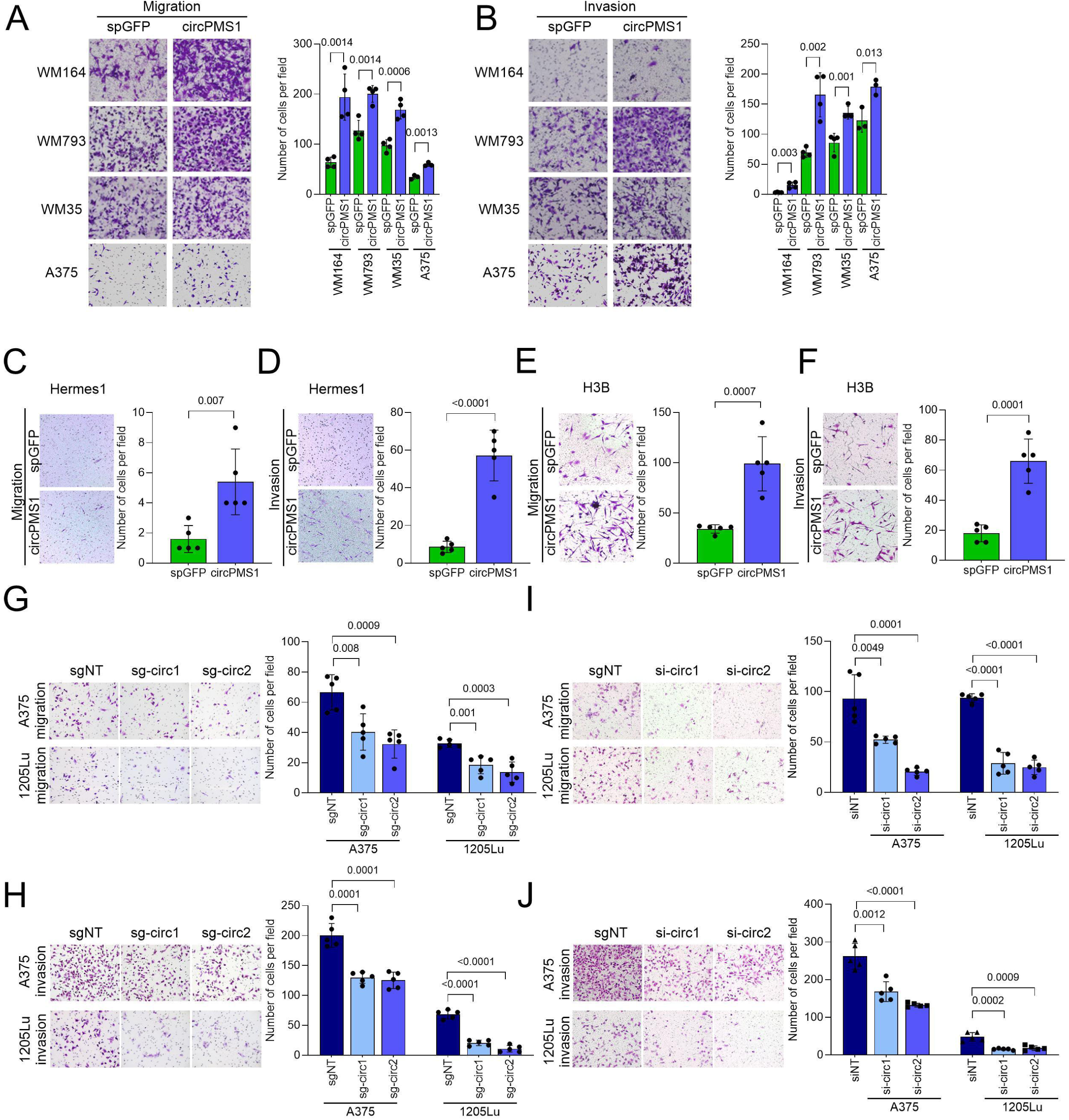
*circPMS1* promotes cell migration and invasion in vitro. (**A**) Transwell migration assay of WM164, WM793, WM35, A375 melanoma cell lines expressing *spGFP* or *circPMS1*. (**B**) Transwell invasion assay of WM164, WM793, WM35, A375 melanoma cell lines expressing *spGFP* or *circPMS1*. (**C**) Transwell migration assay of Hermes1 melanocytes expressing *spGFP* or *circPMS1*. (**D**) Transwell invasion assay of Hermes1 melanocytes expressing *spGFP* or *circPMS1*. (**E**) Transwell migration assay of Hermes3-BRAF^V600E^ melanocytes expressing *spGFP* or *circPMS1*. (**F**) Transwell invasion assay of Hermes3-BRAF^V600E^ melanocytes expressing *spGFP* or *circPMS1*. (**G, H**) Transwell migration (G) and invasion (H) assays of 1205Lu and A375 melanoma cell lines expressing CasRX and either sgNT, sg-circ1, or sg-circ2. (**I, J**) Transwell migration (I) and invasion (J) assays of 1205Lu and A375 melanoma cell lines transfected with siNT, si-circ1, or si-circ2. Statistical significance was determined using t-test. For migration and invasion assays, 4-5 fields were quantified per replicate.

To further validate the functional relevance of *circPMS1*, we performed loss-of-function studies in A375 and 1205Lu melanoma cell lines expressing moderate-to-high endogenous levels of *circPMS1* (**Fig. 1G**). To achieve specificity, we targeted the back-splice junction of *circPMS1* using either CasRX-based RNA editing^42,43^ or siRNA-mediated knockdown to silence the circRNA without also targeting linear *PMS1* (**Supplementary Fig. S2M-P**). *circPMS1* silencing resulted in a significant reduction of both migration and invasion compared with non-targeting controls (**Fig. 2G-J**). These findings demonstrate that *circPMS1* specifically promotes melanoma cell migration and invasion without affecting proliferation.

### *circPMS1* enhances spontaneous melanoma cell dissemination and metastasis

Given the migratory and invasive phenotypes observed *in vitro*, we next determined the effect of *circPMS1* on melanoma metastasis. To this end, we first injected luciferase-labeled WM35 cells overexpressing *circPMS1* or the *spGFP* control via the tail vein into NSG mice. Mice injected with cells expressing ectopic *circPMS1* displayed moderately increased luciferase signal intensity compared with control animals (**Fig. 3A, B**). Histological analysis of organs at endpoint revealed a significant increase in the number metastatic nodules in the lungs, but not the liver (**Fig. 3C-E**). Conversely, *circPMS1* silencing in 1205Lu cells using CasRX followed by tail vein injection in NSG mice significantly reduced luciferase signal intensity as well as the number of metastatic lesions in both lungs and liver (**Fig. 3F-J**). These findings indicate a pro-metastatic role for *circPMS1* in melanoma.

**Fig. 3.**
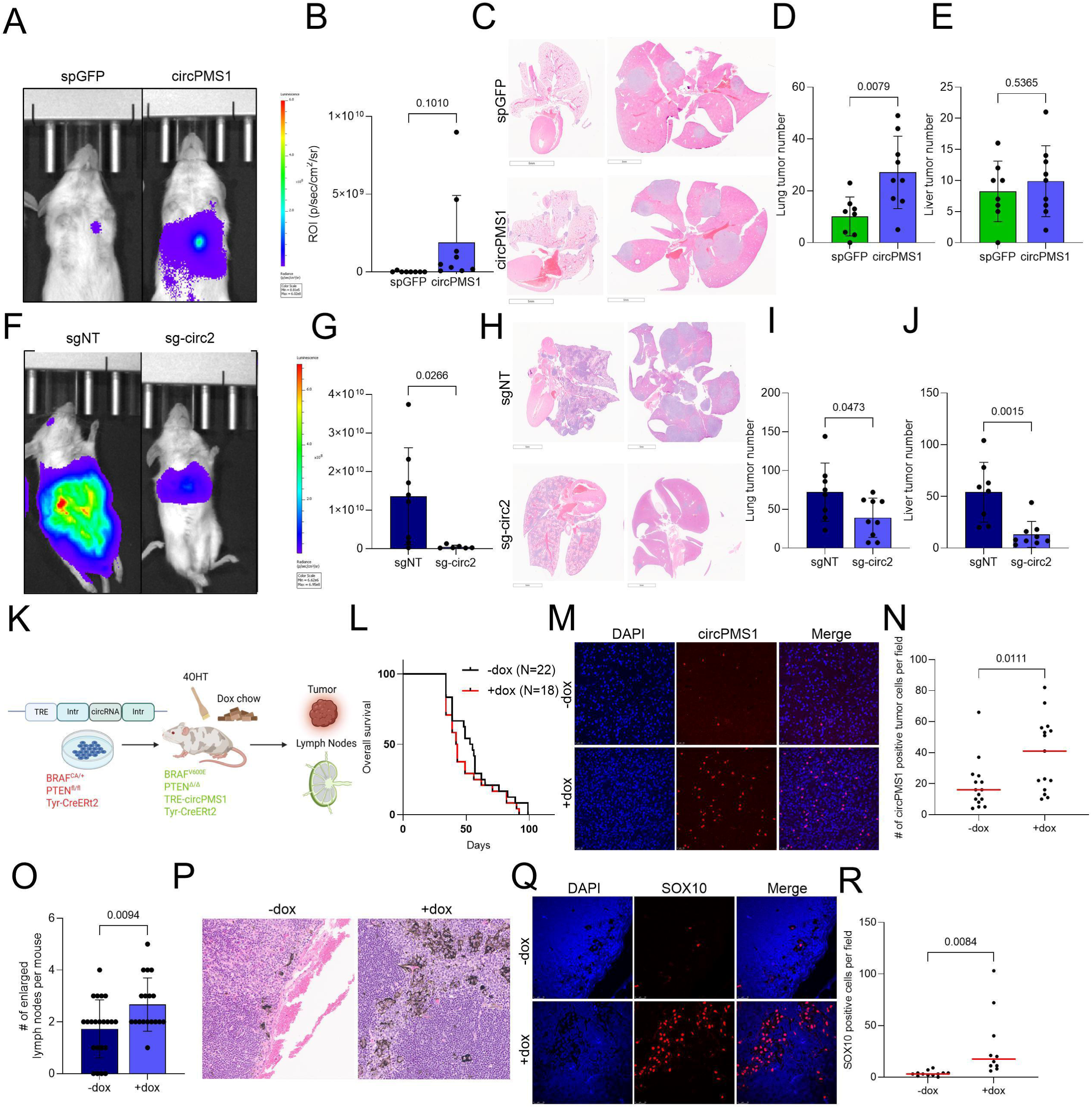
*circPMS1* enhances spontaneous dissemination and metastasis in vivo. (**A**) Bioluminescence imaging of NSG mice injected via the tail vein with WM35 cells expressing *circPMS1* or *spGFP*. (**B**) Quantification of bioluminescence signal in mice shown in (A) (p/sec/cm^2^/sr). n=9 *circPMS1* mice, n=8 *spGFP* mice. (**C**) H&E staining of lungs (left) and livers (right) from NSG mice injected with WM35 *circPMS1* or *spGFP* cells. (**D, E**) Quantification of metastatic nodules in lungs (D) and livers (E) from mice shown in (A-C). (**F**) Bioluminescence imaging of NSG mice injected via the tail vein with 1205Lu cells expressing CasRX and either sgNT or sg-circ2. (**G**) Quantification of bioluminescence signal in mice shown in (F) (p/sec/cm^2^/sr). n=6 sg-circ2 mice, n=8 sgNT mice. (**H**) H&E staining of lungs (left) and livers (right) from NSG mice injected with 1205Lu sgNT or sg-circ2 cells. (**I, J**) Quantification of metastatic nodules in lungs (I) and livers (J) from mice shown in (F-H). (**K**) Scheme of the ES cell-genetically engineered mouse model harboring melanocyte-specific, inducible *circPMS1*, BRAF^V600E^, and PTEN^FL/FL^. (**L**) Kaplan-Meier curve of BPP-circPMS1 mice on regular or dox-containing chow (-dox mice=22, +dox mice=18). (**M**) *circPMS1* fluorescent in situ hybridization (FISH) of tumors from BPP-circPMS1 mice regular or dox-containing chow (n=15 fields per condition). (**N**) Quantification of *circPMS1* FISH show in (M) shown as number of positive cells per field. (**O**) Number of enlarged mandibular and inguinal lymph nodes per BPP-circPMS1 mouse. (**P**) Representative H&E staining of lymph nodes from BPP-circPMS1 mice on either regular or dox chow. (**Q**) SOX10 immunofluorescence staining of lymph nodes from BPP-circPMS1 mice on either regular or dox chow. (**R**) Quantification of SOX10-positive cells per lymph node (n=10 lymph nodes per condition). Statistical significance was determined using t-test.

To investigate the role of *circPMS1* in primary tumor growth and spontaneous dissemination, we used our embryonic stem cell-genetically engineered mouse modeling platform^44^ to generate a mouse model harboring a doxycycline-inducible *circPMS1* transgene on a BRAF^V600E^ mutant and PTEN deleted (Braf^CA^; Pten^FL/FL^; Tyr-CreERt2; Cags-LSL-rtTA3) background (**Fig. 3K**). Three-week-old mice were treated with 4-hydroxytamoxifen (4-OHT) to activate BRAF^V600E^ expression and induce PTEN deletion. Immediately following 4-OHT application, 18 mice were placed on a doxycycline-containing diet to induce *circPMS1* expression (+dox), while 22 control mice were maintained on standard chow (-dox). Animals were monitored for tumor development and sacrificed when tumors reached 2 cm^3^ or became ulcerated, at which point primary tumors and lymph nodes were collected for metastatic assessment (**Fig. 3K**). Primary melanomas formed with complete penetrance and equal latency in +dox and-dox cohorts and, accordingly, no significant differences in overall survival were observed between groups (**Fig. 3L**). This observation combined with the *in vitro* findings support the notion that *circPMS1* does not play a major role in melanoma initiation or primary tumor growth. *circPMS1* expression was validated by FISH, confirming significantly higher *circPMS1* levels in +dox tumors (**Fig. 3M, N**). Upon necropsy, we evaluated mandibular and inguinal lymph nodes and found a significantly higher number of enlarged and pigmented lymph nodes in +dox mice that express ectopic *circPMS1* (**Fig. 3O**). Histological analysis confirmed the increased presence of pigmented cells in the lymph nodes of +dox mice (**Fig. 3P**). Immunofluorescence analysis for the melanoma marker SOX10 demonstrated that pigmentation was attributable to melanoma cell infiltration, with +dox lymph nodes displaying a significantly greater abundance of SOX10-positive cells relative to controls (**Fig. 3Q, R**). While the BRAF^V600E^; PTEN^-/-^ melanoma model may present with micro-metastasis in the lungs, we observed no discernible dissemination to other organs in the-dox and +dox cohorts. These findings indicate that *circPMS1* enhances spontaneous melanoma cell dissemination and lymph node invasion, highlighting a role for *circPMS1* in melanoma progression and metastasis.

### *circPMS1* induces cytoskeletal and focal adhesion remodeling

We next investigated how *circPMS1* promotes the observed pro-metastatic phenotype of melanoma cells. Brightfield examination of melanoma cells with modulated *circPMS1* expression revealed striking changes in cell shape and clustering. Specifically, WM35 melanoma cells overexpressing *circPMS1* adapted a more elongated, mesenchymal-like morphology with extended protrusions and were more dispersed with fewer cells in direct contact with their neighbors (**Fig. 4A**) Conversely, *circPMS1* silencing in A375 cells promoted the formation of compact colonies composed predominantly of rounded cells with fewer protrusions, while control cells retained a more elongated and dispersed phenotype (**Fig. 4A**). These observations suggested that *circPMS1* may regulate cytoskeletal organization and cell adhesion. To further characterize these morphological alterations, WM35 melanoma cells were stained with fluorescent phalloidin to visualize the actin cytoskeleton. *circPMS1* overexpression induced a significantly more elongated cellular morphology compared with control cells (**Fig. 4B, C**). Quantitative analysis of actin fiber structures further demonstrated significant increases in spine area (**Fig. 4D**), length (**Fig. 4E**), and volume (**Fig. 4F**) in *circPMS1* overexpressing cells, indicative of extensive cytoskeletal remodeling. Given the *circPMS1*-driven protrusions and cytoskeletal rearrangements, we tested the effect of *circPMS1* on focal adhesions. To this end, we stained for the focal adhesion-associated protein Paxillin, which plays a central role in mediating cell adhesion, migration and invasion, and communication with the extracellular matrix^54,55^. Consistent with the migratory phenotype, *circPMS1* expression increased the number of Paxillin foci, whereas *circPMS1* silencing resulted in a reduction in Paxillin foci (**Fig. 4G**). Thus, *circPMS1* promotes cytoskeletal and focal adhesion remodeling, driving the acquisition of a more elongated, motile, and invasive cellular phenotype.

**Fig. 4.**
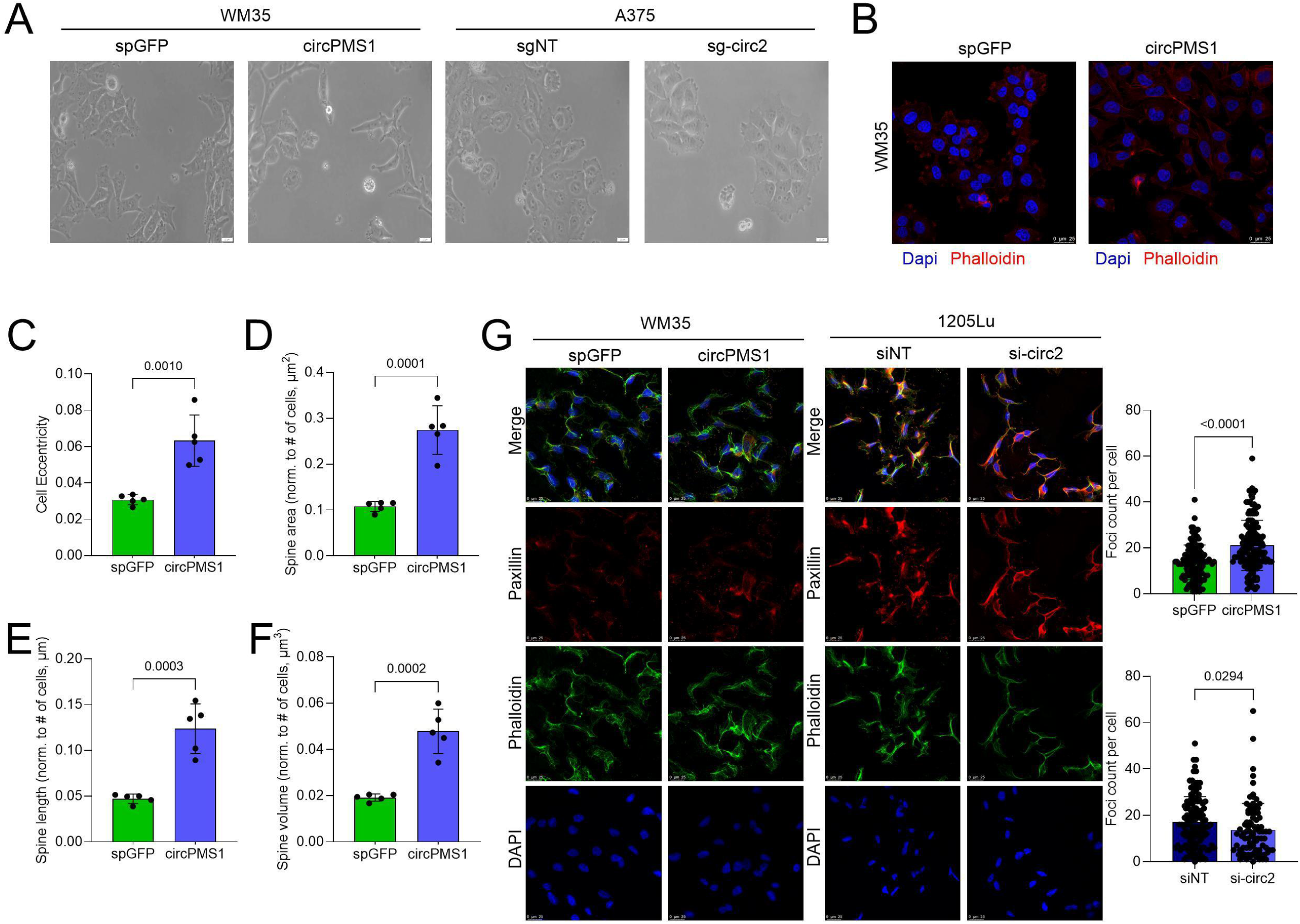
circPMS1 overexpression induces cell shape changes. (**A**) Brightfield images of WM35 cells expressing *spGFP* or *circPMS1* and A375 cells expressing CasRX and either sgNT or sg-circ2. (**B**) Phalloidin immunofluorescence of WM35 cells expressing *spGFP* or *circPMS1*. (**C**) Quantification of cell eccentricity. Greater values indicate a rounder cell shape (n=5 fields per condition). (**D**) Quantification of Phalloidin filament (spine) areas. (**E**) Quantification of Phalloidin filament (spine) lengths. (**F**) Quantification of Phalloidin filament (spine) volumes. Values in (D-F) were normalized to the number of cells per field (n=5 fields per condition). (**G**) Paxillin immunofluorescence in WM35 cells expressing *circPMS1* or *spGFP* (left) and 1205Lu cells transfected with siNT or si-circ2 (middle). Quantifications for WM35 and 1205 cells are shown in top right and bottom right, respectively. Data are plotted as foci per cell. Statistical significance was determined using t-test.

### Circularization, but not translation, is required for the migratory effect of *circPMS1*

While the *ZKSCAN1* introns used in our *circPMS1* expression constructs mediate efficient back-splicing, some ectopic *circPMS1* transcripts may remain linear. To ascertain that the pro-migratory phenotype was specifically mediated by the circular form of *circPMS1* rather than a truncated *PMS1* fragment comprising exons 2-5, we mutated either the upstream splice acceptor (ΔSA) or downstream splice donor (ΔSD) sites required for back-splicing and circularization. We confirmed that the ΔSA or ΔSD mutations effectively abolished *circPMS1* formation (**Supplementary Fig. S3A**), while expression of the mutant constructs was unaffected (**Supplementary Fig. S3B**). Importantly, preventing circularization with either the ΔSA or ΔSD mutant negated the increase in migration and invasion of WM35 melanoma cells elicited by wildtype *circPMS1* (**Supplementary Fig. S3C**). Thus, the metastatic phenotype specifically depends on the circularized form of *PMS1* exons 2-5, rather than a linear, non-circularized transcript containing the same exonic sequence.

Inspecting the *circPMS1* sequence revealed the presence of a putative open reading frame (ORF): *circPMS1* harbors the canonical *PMS1* start codon in exon 2 and the reading frame extends from this ATG through exons 2-5 and across the back-splice junction, which introduces a frameshift that results in a stop codon in the fourth codon. In addition, *in silico* analyses (CircRNADb and circBase) predicted the presence of an Internal Ribosome Entry Site (IRES) within exon 5 of *circPMS1* (**Supplementary Fig. S4A**), suggesting the potential for cap-independent translation of a *circPMS1*-encoded truncated PMS1 protein. To investigate whether an encoded protein mediates the pro-migratory effects of *circPMS1*, we first mutated the ATG start codon in the *circPMS1* overexpression construct (ΔATG). The *circPMS1*-ΔATG mutant was efficiently expressed (**Supplementary Fig. S4B**); however, it failed to enhance migration and invasion of WM793, WM35, and A375 melanoma cells (**Supplementary Fig. S4C-E**), indicating that *circPMS1* could have coding potential. To investigate this further, we first evaluated the predicted IRES sequence using a luciferase reporter assay. In WM793 and WM35 melanoma cells, the putative *circPMS1* IRES enabled translation to an extent similar to its antisense negative control, significantly below the levels observed for a verified IRES from Encephalomyocarditis virus (ECMV) (**Supplementary Fig. S4F**). Next, we generated a *circPMS1* overexpression construct harboring a V5 epitope tag inserted upstream of the first in-frame stop codon within the putative ORF (**Supplementary Fig. S4A**). As a control, we also generated a construct containing the linearized ORF (from the ATG start codon over the back-splice junction to the V5-tag and stop codon; **Supplementary Fig. S4A**) which depends on cap-dependent rather than IRES-dependent translation. Expression of the V5-tagged circular and linear *circPMS1* constructs in HEK293 cells showed that while the linear construct produced a truncated PMS1 protein of the expected size, the circular construct failed to produce a protein (**Supplementary Fig. S4G**). We then assessed whether the protein encoded by the linearized construct could account for the pro-migratory phenotype observed upon *circPMS1* overexpression. Expression of the truncated PMS1 protein via the linearized *circPMS1* construct had no effect on migration and invasion of A375 and WM35 melanoma cells (**Supplementary Fig. S4H**). Collectively, these findings indicate that *circPMS1* is unlikely to be translated and its pro-metastatic effects are therefore not mediated by a cryptic *circPMS1*-encoded truncated PMS1 protein.

### *circPMS1* interacts with LIM-domain proteins ABLIM1, LMO7, and PDLIM7

The functional relevance of the ATG-containing region may reside in its contribution to *circPMS1* secondary structure. Indeed, the ΔATG mutation significantly affects the predicted secondary structure of this region (from RNA Folding Form, https://www.unafold.org/mfold/applications/rna-folding-form.php; **Supplementary Fig. S5A**). Intriguingly, the back-splice junction is located within this motif (**Supplementary Fig. S5A**), providing a possible explanation for the requirement of circularization. RNA secondary structures such as stem loops and double-stranded helices can mediate the interaction with RNA binding proteins (RBPs), and we hypothesized that *circPMS1* elicits its pro-migratory effects through interaction with RBPs.

To identify *circPMS1*-interacting proteins, we created *circPMS1* tagged with the MS2 motif (*MS2-circPMS1*) and confirmed that the *MS2-circPMS1* construct was expressed at levels comparable to untagged *circPMS1* (**Supplementary Fig. S5B**). We then performed RNA pulldown of *MS2-circPMS1* using a MS2-binding protein fused to GST (MBP-GST), with untagged *circPMS1* and pCDNA-MS2 as controls, which yielded an approximately 15-fold enrichment of *MS2-circPMS1* compared to the untagged circRNA (**Fig. 5A**). We expressed and purified *MS2-circPMS1* from both WM35 and A375 cells and subjected the co-precipitated material to mass spectrometry analysis. We identified six associated proteins consistently enriched in the MS2 pulldown over negative controls expressing MS2 loops only (PM/M) or circPMS1 lacking MS2 loops (PM/P): actin-binding LIM protein 1 (ABLIM1), LIM domain only protein 7 (LMO7), PDZ and LIM domain protein 7 (PDLIM7), capping actin protein of muscle Z-line subunit alpha 2 (CAPZA2), coiled-coil-helix-coiled-coil-helix domain containing 6 (CHCHD6), and tropomyosin 1 (TPM1) (**Fig. 5B**). To validate these interacting proteins and prioritize them according to their functional contribution to the *circPMS1*-mediated phenotype, we performed immunoprecipitation followed by RT-qPCR with divergent *circPMS1* primers. Interestingly, ABLIM1, LMO7, and PDLIM1 co-precipitated wildtype *circPMS1* but not ΔATG-mutant *circPMS1* (**Fig. 5C, Supplementary Fig. S5C**), while CAPZA2, CHCHD6, and TPM1 failed to validate in independent pulldown experiments and/or did not show rescue upon ΔATG mutation (**Supplementary Fig. S5D**). These findings suggest that ABLIM1, LMO7, and PDLIM7 interact with *circPMS1* in or near the critical region containing the ATG start codon.

**Fig. 5.**
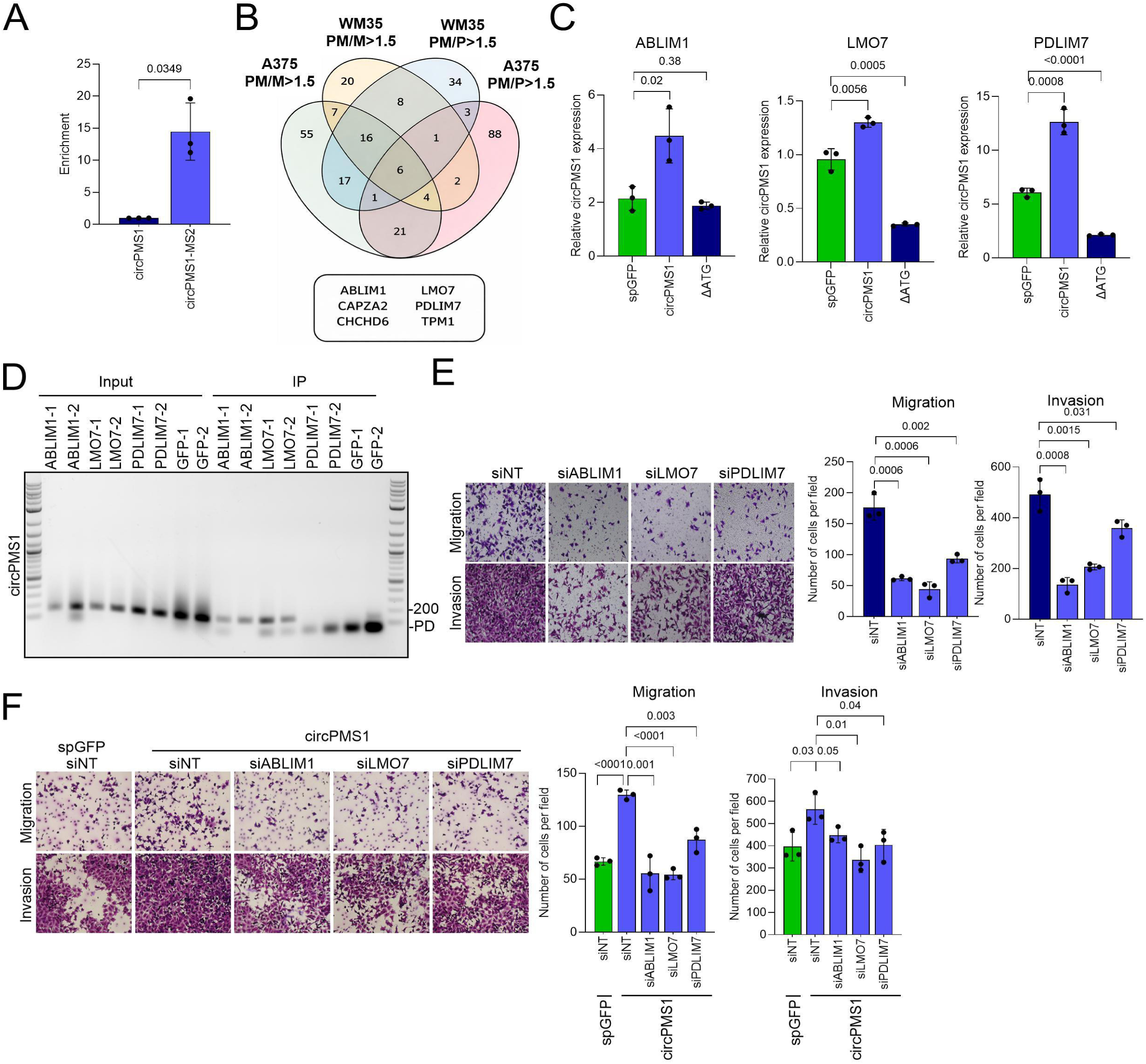
LIM proteins interact with *circPMS1* and mediate its pro-migratory effects. (**A**) Quantification of *circPMS1* enrichment by MS2-mediated pulldown. (**B**) Venn diagram of enriched proteins eluted with circPMS1-MS2 (PM) compared to circPMS1 only (P) and pCDNA3-MS2 (M) controls. (**C**) Quantification of RIP-PCRs for *circPMS1*. ABLIM1, LMO7, or PDLIM7 were immunoprecipitated from HEK293T cells expressing *spGFP*, *circPMS1* or Δ*ATG* and *circPMS1* was detected by RT-qPCR using divergent primers. (**D**) Agarose gel electrophoresis of RT-PCR-amplified *circPMS1* using divergent primers following Covalent Linkage & Affinity Purification (CLAP) of ABLIM1, LMO7, and PDLIM7. PD= primer dimers. (**E**) Transwell migration and invasion assays of WM35 cells after siRNA-mediated silencing of endogenous ABLIM1, LMO7, or PDLIM7. (**F**) Transwell migration and invasion assays of WM35 cells overexpressing *circPMS1* after siRNA-mediated silencing of endogenous ABLIM1, LMO7, or PDLIM7. Statistical significance was determined using t-test.

We next performed crosslinking and affinity purification (CLAP) to determine whether binding to these LIM proteins was direct. We observed direct interaction between *circPMS1* and both ABLIM1 and LMO7, whereas PDLIM7 did not exhibit direct binding (**Fig. 5D**), suggesting that it is recruited as part of a protein complex. Importantly, these interactions were specific to the circular RNA, as no binding was detected with the linear *PMS1* transcript (**Supplementary Fig. S5E**). This further supports the notion that ABLIM1 and LMO7 directly recognize a secondary structure unique to the circular form of *circPMS1*, rather than a primary sequence shared with the linear transcript.

To investigate the functional relevance of these proteins, we assessed the effects of their depletion on the migratory behavior of melanoma cells. Transwell assays revealed that silencing of ABLIM1, LMO7, or PDLIM7 significantly reduced migration and invasion of WM35 cells (**Fig. 5E, Supplementary Fig. S5F, G**) without affecting cell viability (**Supplementary Fig. S5H**). Moreover, silencing ABLIM1, LMO7, or PDLIM7 in the context of *circPMS1* overexpression abrogated the enhanced migratory and invasive phenotype (**Fig. 5F**), indicating that *circPMS1*-mediated phenotypes require the coordinated activity of these LIM domain proteins. These data support a model in which *circPMS1* functions as a structural scaffold that directly engages ABLIM1 and LMO7 and indirectly recruits PDLIM7 to promote melanoma cell migration and invasion.

### circPMS1 scaffolds a complex containing ABLIM1, LMO7 and PDLIM7 at the cytoskeleton

To further elucidate the nature of this functional interaction, we investigated whether *circPMS1* modulates the expression or subcellular localization of the identified LIM proteins. However, no substantial changes were observed in ABLIM1, LMO7, and PDLIM7 expression (**Supplementary Fig. S6A**) or subcellular localization (**Supplementary Fig. S6B**) following *circPMS1* overexpression. We therefore examined whether ABLIM1, LMO7, and PDLIM7 interact with *circPMS1* as part of a multiprotein complex or through mutually exclusive binding mechanisms. To this end, we performed immunoprecipitation assays in WM35 melanoma cells expressing Halo-tagged ABLIM1. Pulldown of ABLIM1 resulted in the co-precipitation of *circPMS1* (**Supplementary Fig. S6C**) as well as LMO7 and PDLIM7 (**Fig. 6A**), suggesting that these three LIM proteins form a complex. To determine whether *circPMS1* is required for complex assembly, lysates were treated with a combination of RNase A and RNase 1 prior to ABLIM1 pulldown. Complete RNA degradation markedly impaired the pulldown efficiency of LMO7 and PDLIM7 (**Fig. 6B**), indicating that *circPMS1* is necessary for scaffolding a complex containing ABLIM1, LMO7, and PDLIM7.

**Fig. 6.**
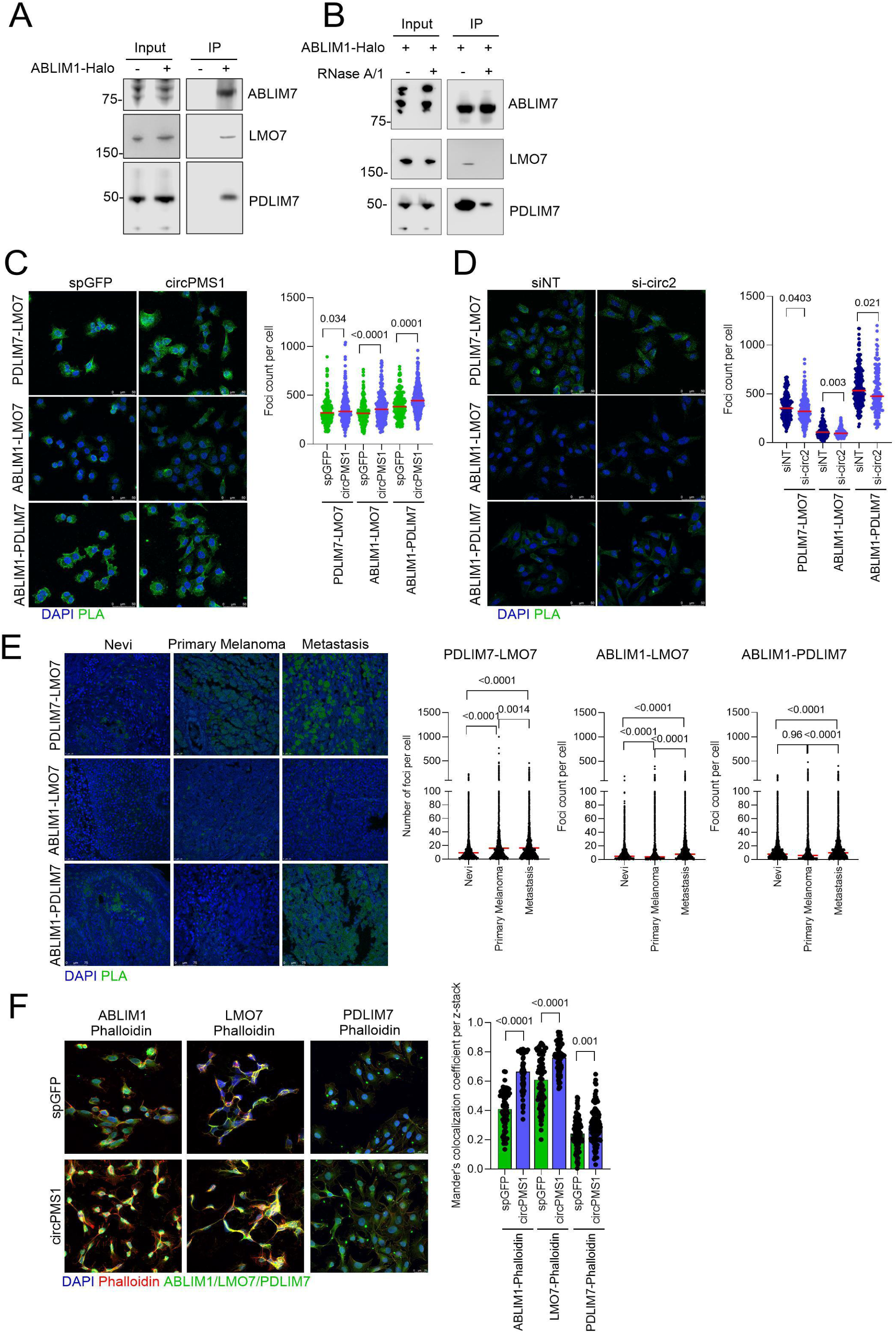
*circPMS1*, ABLIM1, LMO7, and PDLIM7 form a complex. (**A**) Halo-tag pulldown of ABLIM1-Halo from WM35 cells overexpressing *circPMS1*, followed by Western blot for ABLIM1, LMO7, and PDLIM7 (**B**) Halo-tag pulldown and Western blot as in (A), but protein lysates were treated with RNase A/1 cocktail prior to pulldown. (**C**) Proximity ligation assays for PDLIM7-LMO7, ABLIM1-LMO7, or ABLIM1-PDLIM7 in WM35 expressing *circPMS1* or *spGFP*. (**D**) Proximity ligation assay for PDLIM7-LMO7, ABLIM1-LMO7, or ABLIM1-PDLIM7 in 1205Lu cells transfected with siNT or si-circ2. (**E**) Proximity ligation assay for PDLIM7-LMO7, ABLIM1-LMO7, or ABLIM1-PDLIM7 in human melanoma tissue microarrays. Number of foci per cell were analyzed by CellProfiler. (**F**) Co-Immunofluorescence for ABLIM1, LMO7, or PDLIM7 and Phalloidin in WM35 cells expressing *circPMS1* or *spGFP*. Each z-stack field has been analyzed with CellProfiler and is plotted in the graph. Statistical significance was determined using t-test.

We next performed Proximity Ligation Assays (PLA) in melanoma cells to further examine the *circPMS1*-scaffolded protein complex. *circPMS1* overexpression in WM35 cells significantly increased the number of PLA foci representing PDLIM7-LMO7, ABLIM1-LMO7, and ABLIM1-PDLIM7 interactions (**Fig. 6C**). Conversely, *circPMS1* silencing in 1205Lu cells significantly reduced these interactions (**Fig. 6D**). These findings further support *circPMS1*-dependent assembly of the ABLIM1-LMO7-PDLIM7 complex. Consistently, PLA analyses performed on melanoma tissue microarrays (TMAs) revealed a significant enrichment of PLA foci for all three LIM protein pairings in metastatic lesions compared with nevus samples, with some interactions also being elevated in primary tumors (**Fig 6E**). To investigate whether *circPMS1* mediates recruitment of this complex to the cytoskeleton, we performed co-immunofluorescence analyses between each LIM protein and phalloidin. *circPMS1* overexpression increased the co-localization of ABLIM1, LMO7, and PDLIM7 with phalloidin, suggesting enhanced association with actin filaments (**Fig. 6F, Supplementary Fig. S6D**). Collectively, these results support a model in which *circPMS1* promotes the assembly of a protein complex comprising ABLIM1, LMO7, and PDLIM7, thereby facilitating recruitment to actin filaments and promoting cytoskeleton reorganization.

### RhoA inhibition rescues the *circPMS1*-mediated migratory phenotype

To further determine the consequences of *circPMS1* in melanoma cells, we performed RNA sequencing in WM35 melanoma cells overexpressing either *circPMS1* or *spGFP* control, followed by Gene Ontology enrichment analysis (**Fig. 7A**). *circPMS1* overexpression resulted in significant enrichment of pathways associated with cell junction organization, cell migration, and WNT signaling, indicating that *circPMS1*-driven cytoskeletal remodeling promotes secondary activation of pro-migratory signaling programs. Interestingly, virtually all the enriched biological processes identified by RNA sequencing are either directly regulated by or downstream of RhoA signaling^54,56–58^. Although a direct functional interaction between ABLIM1, LMO7, or PDLIM7 and RhoA has not previously been demonstrated, we reasoned that RhoA signaling might nevertheless contribute to the metastatic phenotype induced by *circPMS1*.

**Fig. 7.**
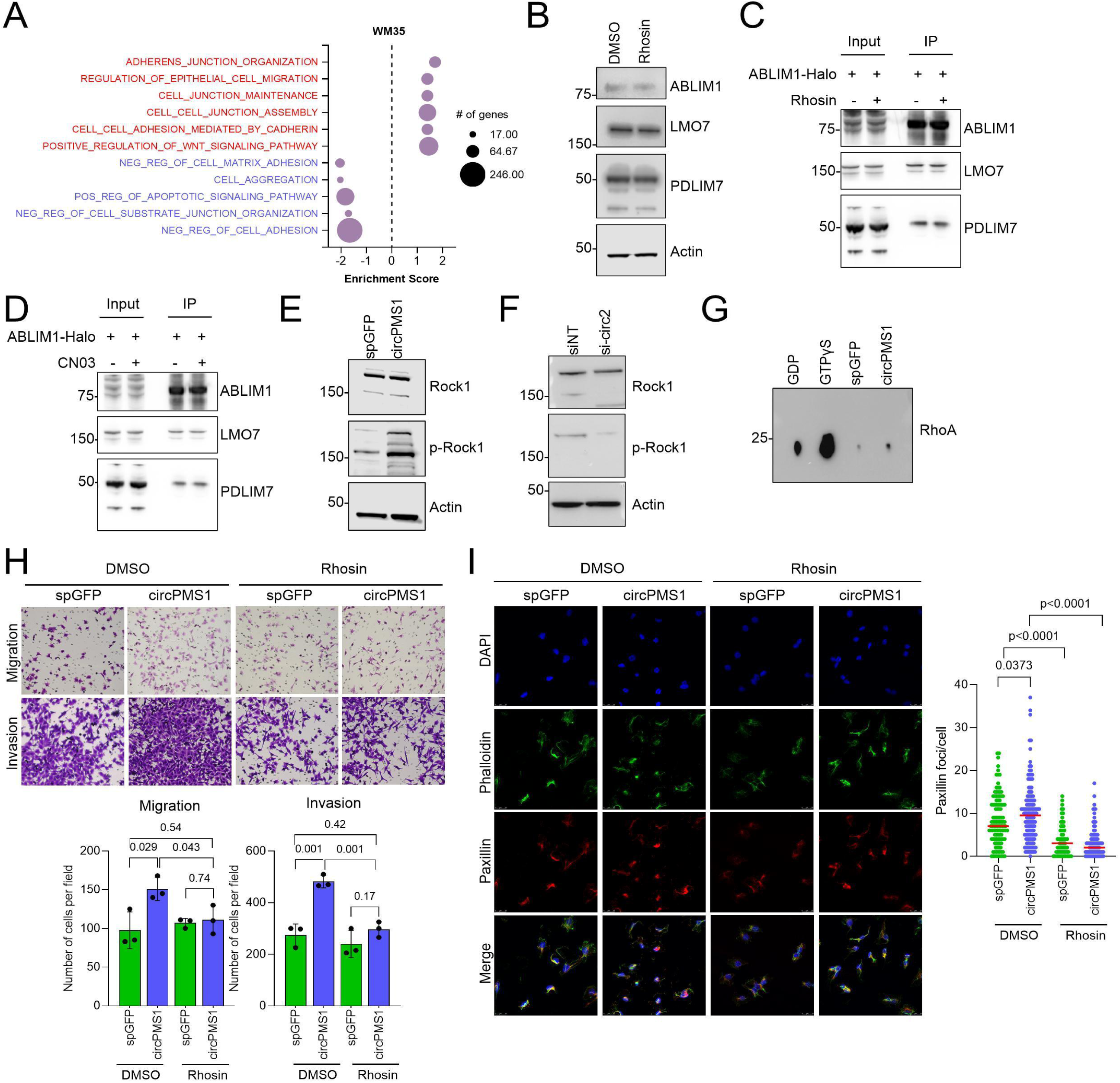
*circPMS1* activates RhoA signaling. (**A**) Gene Ontology analysis on RNA sequencing data obtained from WM35 cells expressing *circPMS1* or *spGFP*. All GO terms shown have p values of 0; size of the circles depicts the number of genes involved in the GO term. (**B**) Western blot for ABLIM1, LMO7, and PDLIM7 in WM35 cells upon RhoA inhibition with Rhosin. Actin was used as a loading control. (**C**) Halo-tag pulldown of ABLIM1-Halo from WM35 cells expressing *circPMS1* and treated with Rhosin or DMSO control, followed by Western blot for ABLIM1, LMO7, and PDLIM7. (**D**) Halo-tag pulldown of ABLIM1-Halo from WM35 cells expressing *circPMS1* and treated with the RhoA activator CN03 or DMSO control, followed by Western blot for ABLIM1, LMO7, and PDLIM7. (**E**) Western blot of p-ROCK1 in WM35 cells expressing *circPMS1* or *spGFP*. Total ROCK1 and Actin were used as loading control. (**F**) Western Blot of p-ROCK1 in WM35 harboring *circPMS1* silencing. (**G**) Western Blot of GTP-bound active RhoA after pulldown. WM35 lysates spiked with GDP and GTPγS are the negative and positive controls, respectively. (**H**) Transwell migration and invasion assays for WM35 cells expressing *circPMS1* or *spGFP* treated with either DMSO or Rhosin. Top, representative images; bottom, quantification. (n = 3 fields). (**I**) Paxillin and Phalloidin immunofluorescence in WM35 cells expressing *circPMS1* or *spGFP* and treated with Rhosin. Foci per cell are plotted. Statistical significance was determined using t-test.

To test whether RhoA signaling is upstream of the *circPMS1*-scaffolded complex, we inhibited the pathway using Rhosin, a selective small-molecule inhibitor of RhoA activation. While Rhosin effectively lowered the levels of the RhoA downstream target p-ROCK1(**Supplementary Fig. S7A**), it had no effect on *circPMS1* expression (**Supplementary Fig. S7B**). In addition, inhibition of RhoA signaling did not affect endogenous expression of ABLIM1, LMO7, or PDLIM7 (**Fig. 7B**). We next investigated whether RhoA activity was required for assembly of the *circPMS1*-scaffolded protein complex. Halo pulldown experiments performed following Rhosin treatment demonstrated that inhibition of RhoA did not disrupt the interaction between ABLIM1, LMO7, and PDLIM7 (**Fig. 7C**). Consistently, pharmacological activation of RhoA using CN03 (**Supplementary Fig. S7C**) similarly had no impact on complex formation (**Fig. 7D**). Thus, RhoA activity does not regulate the expression of *circPMS1* and the LIM proteins and is dispensable for assembly of the *circPMS1*-scaffolded complex.

RhoA signaling is both a driver and a consequence of cytoskeletal reorganization and mechanotransduction, particularly during stress fiber formation and focal adhesion maturation^59,60^. Activation of RhoA may thus occur subsequent to, or be reinforced by, the cytoskeletal remodeling initiated by the *circPMS1*-scaffolded complex. Notably, *circPMS1* overexpression significantly increased p-ROCK1 levels (**Fig. 7E**), whereas *circPMS1* silencing markedly reduces p-ROCK1 levels in WM35 melanoma cells (**Fig. 7F**). Moreover, overexpression of *circPMS1* enhanced the abundance of active, GTP-bound RhoA (**Fig. 7G**), suggesting that RhoA signaling is downstream of the *circPMS1*-scaffolded complex. We then determined whether the pro-migratory effect of *circPMS1* is dependent on RhoA signaling. To this end, WM35 melanoma cells were treated with Rhosin followed by transwell migration and invasion assays. RhoA inhibition had no effect on viability (**Supplementary Fig. S7D**) but significantly impaired migration and invasion induced by *circPMS1* overexpression, whereas no comparable effect was observed in *spGFP* control cells (**Fig. 7F**). Moreover, RhoA inhibition rescued the *circPMS1*-induced Paxillin organization at focal adhesions (**Fig. 7G**), demonstrating that RhoA signaling mediates the effects of *circPMS1* on the cytoskeletal remodeling and migratory behavior of melanoma cells.

## Discussion

Non-mutational mechanisms, including the dysregulation of non-coding RNAs, are increasingly recognized as drivers of melanoma progression. Here we identify and characterize *circPMS1*, a circular RNA that promotes melanoma cell migration, invasion, and metastatic dissemination. We find that *circPMS1* acts as an RNA scaffold that directly engages the LIM-domain proteins ABLIM1 and LMO7 and recruits PDLIM7 into a cytoskeleton-associated complex, thereby enhancing actin reorganization, focal adhesion dynamics, and cell motility. These findings define a circRNA-based mechanism that links RNA scaffolding to the cytoskeletal machinery of metastasis and expand the understanding of circRNA function in melanoma.

We identified *circPMS1* using a strategy that prioritized circRNAs not only by differential abundance but also by their circular-to-linear splicing ratio, reasoning that circRNAs whose relative circularization is altered in melanoma are more likely to be regulated, functional species rather than by-products of parental gene transcription. *circPMS1* is conserved between human and mouse, expressed across a range of normal murine tissues, and detectable in multiple cancer lineages. Notably, the parental *PMS1* gene encodes a DNA mismatch repair component and is therefore generally regarded as tumor-suppressive^61,62^, yet the circular product of the same locus is oncogenic and pro-metastatic. *circPMS1* thus represents an instance in which a single gene gives rise to two products with opposing roles in cancer, and its activity is independent of the mismatch repair function of linear *PMS1*.

Functionally, *circPMS1* acts as a driver of melanoma cell motility. Gain-and loss-of-function studies in human and murine melanoma cells showed that *circPMS1* enhances migration and invasion independently of oncogenic driver and species context, while leaving proliferation and low-density focus formation unaffected. Consistent with these in vitro phenotypes, *circPMS1* promoted metastasis in vivo. In xenograft assays its overexpression increased metastatic burden while its silencing suppressed dissemination, and in an inducible BRAF^V600E^; PTEN^null^ mouse model, *circPMS1* did not alter primary tumor onset, penetrance, or growth but markedly increased spontaneous dissemination and lymph node metastasis. Together, these data indicate that *circPMS1* selectively promotes the metastatic stages of melanoma rather than tumor initiation or primary growth.

This stage-restricted activity fits an emerging pattern in the melanoma circRNA literature. The most rigorously characterized example, *CDR1as*, is downregulated in melanoma, and its loss drives invasion and metastatic progression rather than tumor initiation^63^. Moreover, *circZNF609* is downregulated in melanoma, where its loss promotes migration and metastasis^64^, while *circ-GLI1* upregulation promotes metastasis and angiogenesis^65^, and several miRNA sponge circRNAs have been reported to enhance migration and invasion of melanoma cells^66^. Although many of these circRNAs remain superficially characterized, their recurrent engagement of migratory, invasive, and metastatic phenotypes supports the notion that circRNA dysregulation in melanoma may contribute preferentially to the later stages of melanomagenesis.

A notable strength of this study is the set of genetic tools that enabled stable *circPMS1* expression in both cells and mice. The gain-of-function experiments reported here rely on a transposon-based system we previously developed for durable circRNA overexpression in cultured cells^41^, which we extended to the autochthonous setting by integrating a doxycycline-inducible *circPMS1* transgene into a BRAF^V600E^; PTEN^null^ background using our embryonic stem cell-based modeling platform^44^. This generated a GEMM for melanocyte-specific overexpression of a naturally occurring circRNA. Because the model preserves tumor-microenvironment interactions and recapitulates the stepwise progression of the disease, it allowed us to attribute *circPMS1* activity specifically to dissemination and lymph node metastasis within the full course of tumor evolution.

circRNAs can function through several mechanisms, including sequestration of miRNAs, regulation of transcription and splicing, templating of translated peptides, and scaffolding of proteins^14–29,31–38,66^. Two lines of evidence indicate a scaffolding function for *circPMS1* and argue against a coding function. First, the pro-migratory activity required circularization, as constructs that could not back-splice failed to enhance migration and invasion despite producing the same exonic sequence in linear form. Second, although *circPMS1* contains a putative ORF and a predicted IRES, we detected no translation from the circle and the truncated *PMS1* protein produced from a linearized construct had no effect on motility, indicating that the phenotype is not mediated by an encoded peptide. The requirement for circularization therefore reflects a structural property of the circle rather than its coding potential. Consistent with this, the start codon lies within a region predicted to form a structured element spanning the back-splice junction, and its mutation abolished activity, implicating this region as a putative structural determinant of *circPMS1* function.

We found that *circPMS1* binds ABLIM1 and LMO7 directly and recruits PDLIM7 into the resulting complex. This binding is specific to the circular form, as neither protein associated with linear *PMS1* nor mutation of the start codon within the putative structured element abolished binding. ABLIM1 and LMO7 therefore appear to recognize a structural feature unique to the circle rather than a sequence shared with the linear transcript. LIM-domain proteins have not generally been recognized as direct RNA binders, and although LIM domains are zinc-finger modules structurally related to nucleic-acid-binding folds, they have been characterized almost exclusively as mediators of protein-protein interactions in cytoskeletal and transcriptional settings^67–78^. Our data indicate that at least some members of this family can bind RNA directly, broadening their known functional repertoire. *circPMS1* thus adds to a still small set of melanoma circRNAs shown to act through RNA-binding proteins, alongside *CDR1as* (IGF2BP3)^63^, *circZNF609* (FMRP)^64^, and *circFCHO2* (DND1)^79^, though many such circRNA-RBP interactions are well documented in normal tissues and other cancers^11^. Within melanoma, *circPMS1* is distinguished by scaffolding cytoskeletal LIM-domain proteins and thereby linking circRNA activity to actin regulation. A related link is seen with *circZNF609*, which suppresses melanoma migration and metastasis through FMRP-mediated destabilization of the Rho-family GTPase RAC1^64^, placing a second melanoma circRNA at the interface of RBP interaction and cytoskeletal control, though *circPMS1* acts in the opposite direction and through a distinct, scaffold-based mechanism.

ABLIM1, LMO7, and PDLIM7 are actin-associated proteins with established roles in cytoskeletal organization, cell adhesion, and motility^77,78,80,81^, and their coordinated recruitment by *circPMS1* provides a direct route from a circRNA-scaffolded complex to the migratory phenotype. The three proteins associate with one another in a *circPMS1*-dependent manner, as degradation of RNA disrupted their co-precipitation and modulating *circPMS1* levels changed the frequency of their interactions in situ, indicating that the circRNA holds the complex together rather than simply binding its individual members. *circPMS1* also increased the association of all three proteins with actin filaments, consistent with the circRNA directing the assembled complex to the cytoskeleton where it promotes remodeling. Notably, these interactions were enriched in metastatic melanoma lesions relative to benign nevi, linking *circPMS1*-dependent complex assembly to disease progression in patient samples and reinforcing its relevance to the metastatic phenotype.

Finally, our data connect the *circPMS1* scaffold to RhoA, a central regulator of actin dynamics^56,58^. *circPMS1* increased the levels of active RhoA signaling, and its inhibition with Rhosin reversed the migratory and invasive phenotype, identifying RhoA as a required effector of *circPMS1*. Manipulating RhoA activity in either direction did not alter assembly of the ABLIM1-LMO7-PDLIM7 complex, indicating that RhoA does not control scaffold formation but instead acts downstream of it. We therefore favor a model in which *circPMS1* nucleates the LIM-domain protein complex on actin filaments, and the resulting cytoskeletal and focal adhesion remodeling activates RhoA, a well-documented feedback relationship^54,57,82^, which in turn reinforces actin reorganization and sustains the motile, invasive state. In this model RhoA operates as a downstream amplifier of a structural event initiated by the circRNA scaffold, linking a discrete RNA-protein interaction to a broadly acting cytoskeletal signaling axis.

Together, these findings define *circPMS1* as a structural scaffold that assembles ABLIM1, LMO7, and PDLIM7 on the actin cytoskeleton to drive the remodeling and RhoA-dependent signaling that underlie melanoma cell motility and metastasis. This work extends the functional repertoire of circRNAs beyond their established roles in RNA and transcriptional regulation to the direct control of cytoskeletal architecture, and it identifies LIM-domain proteins as a previously unappreciated class of RNA-binding partners. Because *circPMS1* acts selectively on dissemination rather than tumor formation, the *circPMS1*-LIM protein axis represents a potential point of intervention for limiting metastatic progression, which remains the main cause of mortality in melanoma. More broadly, our results raise the possibility that structured circRNAs act as organizing centers that couple RNA-protein interactions to the actin cytoskeleton to govern cell movement, a mode of regulation that is unlikely to be confined to melanoma.

## ACKNOWLEDGEMENTS

We thank members of the Karreth lab for helpful discussions and Pietro Berico for critical reading of the manuscript. N.M. acknowledges support from an NCI Predoctoral to Postdoctoral Fellow Transition Award (F99CA305524). O.V.P. was supported by Instituto de Salud Carlos III and co-funded by the European Union under Grants FORT23/0006, PI24/00291, and CP24/00005. X.X. received support from a Miles for Moffitt Postdoc Award and a Melanoma Research Foundation Career Development Award (1068914). F.A.K. was supported by NIH grants R01CA259046 and R21CA299163. This work was also supported by the Gene Targeting Core, Bioinformatics and Biostatistics Shared Resource, Molecular Genomics Core, Analytical Microscopy Core, and Proteomics and Metabolomics Core, which are funded in part by Moffitt’s Cancer Center Support Grant (P30CA076292).

## AUTHOR CONTRIBUTIONS STATEMENT

N.M., O.V., and F.A.K. conceived the project and N.M. and F.A.K. designed experiments. N.M. performed experiments and O.V., M.C., X.X., K.W., M.M., A.A.N., and N.P. assisted with experiments. K.M.G provided assistance with microscopy analyses. M.A.P. and X.Y. performed circRNA identification. J.Y. and X.Y. performed RNA-seq analysis. A.M.J. performed proteomics analysis. F.A.K. supervised the studies and acquired funding.

## COMPETING INTERESTS STATEMENT

The authors declare no competing interests.

## Supplementary Figure Legends

**Sup. Fig. 1.** *circPMS1* characterization across species and cell types.

(**A**) Relative circularization of *circPMS1* (calculated as *circPMS1*/(*circPMS1*+*PMS1*)) in human melanoma cell lines. Each dot represents an RNA-seq replicate. (**B**) Scheme of primer positions in *circPMS1* and *PMS1*. Dark blue primers are divergent, and light blue primers are convergent. (**C**) Sanger sequencing confirming the back-splice junction of human *circPMS1*. (**D**) RT-qPCR for *PMS1* expression in a panel of melanocytes (dark blue), melanocytes harboring BRAF^V600E^ expression (medium blue), and melanoma cell lines (light blue). (**E**) RT-qPCR for endogenous *circPMS1* expression in pancreatic cancer (T3M4 and PATU8988), ovarian cancer (OVC5 and OVC8) and lung adenocarcinoma (A549) cell lines. Expression is shown relative to WM35 melanoma cells. (**F**) Agarose gel confirming the expected size of mouse *circPMS1* back-splice junction PCR using divergent primers. (**G**) Sanger sequencing confirming the back-splice junction of mouse *circPMS1*. (**H**) RT-qPCR of mouse *circPMS1* in different tissues and organs. Expression is shown relative to skin. (**I**) *circPMS1* fluorescent in situ hybridization (FISH) on WM35 cells overexpressing *circPMS1* or *spGFP*. Representative images (left) and quantification in cytoplasm and nuclei (right) are shown (n= 5 fields). Statistical significance was determined using t-test.

**Sup. Fig. 2. *circPMS1* enhances the migratory and invasive potential of cells in vitro.**

(**A**) Confirmation of *circPMS1* overexpression by RT-qPCR in WM164, WM793, WM35, A375 cells. (**C-**E) Proliferation assays of WM164 (B), WM793 (C), WM35 (D), and A375 (E) cells overexpressing *circPMS1* or *spGFP* (n = 3 fields). (**F, G**) Focus formation assay of WM164 and WM793 cells (F) and A375 and WM35 cells (G) overexpressing *circPMS1* or *spGFP* (n = fields). Representative images (bottom) and quantification (top) are shown. (**H, I**) Confirmation of *circPMS1* overexpression by RT-qPCR in Hermes1 (H) and Hermes3-Braf^V600E^ (I) cells. (**J**) Confirmation of human *circPMS1* overexpression by qRT-PCR in murine NCC, NPP, BCC, BPP melanoma cell lines. (**K, L**) Transwell migration (K) and invasion (L) assays in mouse melanoma cell lines overexpressing *circPMS1* or *spGFP* (n= 5 fields). Representative images (left) and quantification (right) are shown. (**M, N**) RT-qPCR quantifying the effect of CasRX-mediated silencing on endogenous *circPMS1* (M) and *PMS1* (N) expression in A375 and 1205Lu cells. (**O, P**) RT-qPCR quantifying the effect of siRNA-mediated silencing on endogenous *circPMS1* (O) and *PMS1* (P) expression in A375 and 1205Lu cells. Statistical significance was determined using t-test.

**Sup. Fig. 3. *circPMS1* circularization drives the pr-migratory phenotype.**

(A) Agarose gel electrophoresis of cDNA amplified using divergent primers to detect circularization of wildtype, ΔSD, and ΔSA *circPMS1* expressed in HEK293T cells. (B) Agarose gel electrophoresis of cDNA amplified using convergent primers to detect expression of wildtype, ΔSD, and ΔSA *circPMS1* expressed in HEK293T cells. (C) Transwell migration and invasion assays in WM35 cells expressing wildtype, ΔSD, or ΔSA circPMS1, or spGFP (n = 3 fields). Representative images (top) and quantification (bottom) are shown. Statistical significance was determined using t-test.

**Sup. Fig. 4. *circPMS1* does not promote migration and invasion via an encoded protein.**

(A) Top: scheme for *circPMS1* overexpression construct containing a V5 tag cloned upstream of start codon to express circularized *circPMS1*. The putative IRES in exon 5 is indicated. Bottom: scheme of the linearized putative *circPMS1*-encoded ORF (cod_circ-v5) in a lentiviral construct.

(B) Confirmation of ectopic expression of the ΔATG *circPMS1* construct relative to ectopic wildtype *circPMS1* expression. (**C-E**) Transwell migration and invasion assays in WM793 (C), WM35 (D), and A375 (E) cells expression wildtype or ΔATG *circPMS1* or *spGFP* (n = 3 fields). Representative images (top) and quantification (bottom) are shown. (**F**) Luciferase assay testing IRES functionality. The predicted *circPMS1* IRES sequence (IRES) is compared to antisense circPMS1 IRES (As_IRES) as negative control and EMCV IRES as positive control. (**G**) Western blot for V5-tagged *circPMS1*-encoded truncated PMS1 protein in HEK293T cells. Truncated PMS1 is detectable when expressed from linearized cod_circ-v5 but not from circularized circPMS1-v5. (**H**) Transwell migration and invasion assays in A375 and WM35 cells comparing the truncated PMS1 protein encoded by cod_circ-v5 and GFP control (n = 5 fields). Representative images (top) and quantification (bottom) are shown. Statistical significance was determined using t-test.

**Sup. Fig. 5. circPMS1 scaffolds an ABLIM1-LMO7-PDLIM7 complex.**

(**A**) Secondary structure prediction of wildtype and ΔATG *circPMS1*. Red squares indicate the wildtype and mutated start codon, BSJ indicated the back-splice junction. (**B**) RT-qPCR confirming efficient expression of the circPMS1-MS2 construct compared to wildtype *circPMS1*.

(C) Western blots to confirm immunoprecipitation of ABLIM1, LMO7, and PDLIM7. (**D**) Quantification of RIP-PCRs for *circPMS1*. CAPZA2, CHCHD6, or TPM1 were immunoprecipitated from HEK293T cells expressing *spGFP*, *circPMS1* or Δ*ATG* and *circPMS1* was detected by RT-qPCR using divergent primers. (**E**) CLAP followed by agarose gel electrophoresis for linear *PMS1* after immunoprecipitation of ABLIM1, LMO7, or PDLIM7. (**F**) Western Blots for ABLIM1, LMO7, or PDLIM7 to confirm siRNA-mediated silencing in WM35 cells. (**G**) RT-qPCR to confirm effective silencing of ABLIM1, LMO7 and PDLIM7 transcripts by siRNAs. (**H**) SytoxGreen cell death assay performed in WM35 cells after ABLIM1, LMO7, or PDLIM7 siRNA transfection. Cell death was analyzed 36 hours post-transfection. Statistical significance was determined using t-test.

**Sup. Fig. 6. ABLIM1-LMO7-PDLIM7 complex formation.**

(**A**) Western Blot for endogenous ABLIM1, LMO7, or PDLIM7 expression upon *circPMS1* overexpression. (**B**) Immunofluorescence for ABLIM1, LMO7, or PDLIM7 in WM35 cells expressing *circPMS1* or *spGFP*. No overt effects on protein localization upon *circPMS1* expression were evident. (**C**) *circPMS1* detection by RT-qPCR following ABLIM1-Halo pulldown. PD=primer dimers. (**D**) Co-Immunofluorescence of Phalloidin with ABLIM1, LMO7, or PDLIM7 in WM35 cells expressing *circPMS1* or *spGFP*.

**Sup. Fig. 7. The effect of RhoA inhibition on cell death and circPMS1 expression.**

(**A**) Western blot of p-ROCK1 upon Rhosin treatment in WM35 cells expressing *circPMS1* or *spGFP*. Total ROCK1 and Actin were used as loading controls. (**B**) RT-qPCR for *circPMS1* upon DMSO or Rhosin treatment in WM35 cells expressing *circPMS1* or *spGFP*. (**C**) Western blot of p-ROCK1 upon CN03 treatment in WM35 cells expressing *circPMS1* or *spGFP*. Total ROCK1 and Actin were used as loading controls. (**D**) SytoxGreen staining of WM35 cells expressing *spGFP* or *circPMS1* treated with either DMSO or Rhosin. Cell death was analyzed 36 hours post-treatment. Statistical significance was determined using t-test.

## Notes

### Competing Interest Statement

The authors have declared no competing interest.

## References

1. Alexandrov, L. B., et al. Signatures of mutational processes in human cancer. Nature 500, 415–421 (2013).

2. Hayward, N. K., et al. Whole-genome landscapes of major melanoma subtypes. Nature 545, 175–180 (2017).

3. Shain, A. H., et al. Genomic and Transcriptomic Analysis Reveals Incremental Disruption of Key Signaling Pathways during Melanoma Evolution. Cancer Cell 34, 45–55.e4 (2018).

4. Shain, A. H., et al. The Genetic Evolution of Melanoma from Precursor Lesions. New England Journal of Medicine 373, 1926–1936 (2015).

5. Akbani, R., et al. Genomic Classification of Cutaneous Melanoma. Cell 161, 1681–1696 (2015).

6. Turner, N., Ware, O. & Bosenberg, M. Genetics of metastasis: melanoma and other cancers. Clin. Exp. Metastasis 35, 379–391 (2018).

7. Vergara, I. A., et al. Evolution of late-stage metastatic melanoma is dominated by aneuploidy and whole genome doubling. Nat. Commun. 12, (2021).

8. Rambow, F., Marine, J.-C. & Goding, C. R. Melanoma plasticity and phenotypic diversity: therapeutic barriers and opportunities. Genes & Dev. 33, 1295–1318 (2019)

9. Sarkar, D., Leung, E. Y., Baguley, B. C., Finlay, G. J. & Askarian-Amiri, M. E. Epigenetic regulation in human melanoma: Past and future. Epigenetics vol. 10 103–121 (2015).

10. Vera, O., Jasani, N. & Karreth, F. A. Long Non-Coding RNAs in Melanoma Development and Biology. Proceedings of the Singapore National Academy of Science vol. 14, 145–166 (2020).

11. Chen, L. L. The expanding regulatory mechanisms and cellular functions of circular RNAs. Nature Reviews Molecular Cell Biology vol. 21, 475–490 (2020).

12. Xiao, M. S., Ai, Y. & Wilusz, J. E. Biogenesis and Functions of Circular RNAs Come into Focus. Trends in Cell Biology vol. 30, 226–240 (2020).

13. Chen, L. L. & Yang, L. Regulation of circRNA biogenesis. RNA Biol. 12, 381–388 (2015).

14. Memczak, S., et al. Circular RNAs are a large class of animal RNAs with regulatory potency. Nature 495, 333–338 (2013).

15. Hansen, T. B., et al. Natural RNA circles function as efficient microRNA sponges. Nature 495, 384–388 (2013).

16. Capel, B., et al. Circular transcripts of the testis-determining gene Sry in adult mouse testis. Cell 73, 1019–1030 (1993).

17. Salmena, L., Poliseno, L., Tay, Y., Kats, L. & Pandolfi, P. P. A ceRNA hypothesis: The rosetta stone of a hidden RNA language? Cell vol. 146, 353–358 (2011).

18. Karreth, F. A. & Pandolfi, P. P. CeRNA cross-talk in cancer: When ce-bling rivalries go awry. Cancer Discov. 3, 1113–1121 (2013).

19. Du, W. W., et al. Induction of tumor apoptosis through a circular RNA enhancing Foxo3 activity. Cell Death Differ. 24, 357–370 (2017).

20. Du, W. W., et al. Foxo3 circular RNA retards cell cycle progression via forming ternary complexes with p21 and CDK2. Nucleic Acids Res. 44, 2846–2858 (2016).

21. Holdt, L. M., et al. Circular non-coding RNA ANRIL modulates ribosomal RNA maturation and atherosclerosis in humans. Nat. Commun. 7, (2016).

22. Abdelmohsen, K., et al. Identification of HuR target circular RNAs uncovers suppression of PABPN1 translation by CircPABPN1. RNA Biol. 14, 361–369 (2017).

23. Yang, Q., et al. A circular RNA promotes tumorigenesis by inducing c-myc nuclear translocation. Cell Death Differ. 24, 1609–1620 (2017).

24. Yang, Z. G., et al. The Circular RNA Interacts with STAT3, Increasing Its Nuclear Translocation and Wound Repair by Modulating Dnmt3a and miR-17 Function. Molecular Therapy 25, 2062–2074 (2017).

25. Li, Z., et al. Exon-intron circular RNAs regulate transcription in the nucleus. Nat. Struct. Mol. Biol. 22, 256–264 (2015).

26. Zhang, Y., et al. Circular Intronic Long Noncoding RNAs. Mol. Cell 51, 792–806 (2013).

27. Conn, V. M., et al. A circRNA from SEPALLATA3 regulates splicing of its cognate mRNA through R-loop formation. Nat. Plants 3, (2017).

28. Wang, Y. & Wang, Z. Efficient backsplicing produces translatable circular mRNAs. RNA 21, 172–179 (2015).

29. Pamudurti, N. R. et al. Translation of CircRNAs. Mol. Cell 66, 9–21.e7 (2017).

30. Yang, Y., et al. Extensive translation of circular RNAs driven by N 6-methyladenosine. Cell Res. 27, 626–641 (2017).

31. Begum, S., Yiu, A., Stebbing, J. & Castellano, L. Novel tumour suppressive protein encoded by circular RNA, circ-SHPRH, in glioblastomas. Oncogene vol. 37, 4055–4057 (2018).

32. Zhang, M., et al. A novel protein encoded by the circular form of the SHPRH gene suppresses glioma tumorigenesis. Oncogene 37, 1805–1814 (2018).

33. Liang, W. C., et al. Translation of the circular RNA circβ-catenin promotes liver cancer cell growth through activation of the Wnt pathway. Genome Biol. 20, (2019).

34. Yang, Y., et al. Novel Role of FBXW7 Circular RNA in Repressing Glioma Tumorigenesis. J. Natl. Cancer Inst. 110, (2018).

35. Zheng, X., et al. A novel protein encoded by a circular RNA circPPP1R12A promotes tumor pathogenesis and metastasis of colon cancer via Hippo-YAP signaling. Mol. Cancer 18, (2019).

36. Zhang, M., et al. A peptide encoded by circular form of LINC-PINT suppresses oncogenic transcriptional elongation in glioblastoma. Nat. Commun. 9, (2018).

37. Legnini, I., et al. Circ-ZNF609 Is a Circular RNA that Can Be Translated and Functions in Myogenesis. Mol. Cell 66, 22–37.e9 (2017).

38. Ho-Xuan, H., et al. Comprehensive analysis of translation from overexpressed circular RNAs reveals pervasive translation from linear transcripts. Nucleic Acids Res. 48, 10368–10382 (2020).

39. Vera, O., et al. A MAPK/miR-29 axis suppresses melanoma by targeting MAFG and MYBL2. Cancers (Basel). 13, 1–18 (2021).

40. Bok, I., Angarita, A., Douglass, S. M., Weeraratna, A. T. & Karreth, F. A. A Series of BRAF-and NRAS-Driven Murine Melanoma Cell Lines with Inducible Gene Modulation Capabilities. JID Innov. 2, (2022).

41. Mecozzi, N., et al. Genetic tools for the stable overexpression of circular RNAs. RNA Biol. 19, 353–363 (2022).

42. Zhang, Y., et al. Optimized RNA-targeting CRISPR/Cas13d technology outperforms shRNA in identifying functional circRNAs. Genome Biol. 22, (2021).

43. Konermann, S., et al. Transcriptome Engineering with RNA-Targeting Type VI-D CRISPR Effectors. Cell 173, 665–676.e14 (2018).

44. Bok, I., et al. A versatile ES cell-based melanoma mouse modeling platform. Cancer Res. 80, 912–921 (2020).

45. Schnack Nielsen Editor, B. In Situ Hybridization Protocols. Springer *US* 2148, (2020).

46. Wu, W., Luo, B., Xiao, Z. An optimized protocol for whole mount in situ hybridization of mouse brain. Int J Clin Med 11(3), 2313–2318 (2018).

47. Dobin, A., et al. STAR: Ultrafast universal RNA-seq aligner. Bioinformatics 29, 15–21 (2013).

48. Zhang, X. O., et al. Diverse alternative back-splicing and alternative splicing landscape of circular RNAs. Genome Res. 26, 1277–1287 (2016).

49. Love, M. I., Huber, W. & Anders, S. Moderated estimation of fold change and dispersion for RNA-seq data with DESeq2. Genome Biol. 15, (2014).

50. Cheng, J., Metge, F. & Dieterich, C. Specific identification and quantification of circular RNAs from sequencing data. Bioinformatics 32, 1094–1096 (2016).

51. Martin, M. Cutadapt Removes Adapter Sequences from High-Throughput Sequencing Reads. EMBnet Journal, 17, 10–12 (2011)

52. Li, B. & Dewey, C. N. RSEM: Accurate transcript quantification from RNA-Seq data with or without a reference genome. BMC Bioinformatics 12, (2011).

53. Bushnell, B., Rood, J. & Singer, E. BBMerge – Accurate paired shotgun read merging via overlap. PLoS One 12, e0185056 (2017).

54. Turner, C. E. Paxillin and Focal Adhesion Signalling. Nat Cell Biol vol. 2 (2000).

55. Ripamonti, M., Wehrle-Haller, B. & de Curtis, I. Paxillin: A Hub for Mechano-Transduction from the β3 Integrin-Talin-Kindlin Axis. Frontiers in Cell and Developmental Biology vol. 10 (2022).

56. Sahai, E. & Marshall, C. J. RHO - GTPases and cancer. Nature Reviews Cancer vol. 2, 133–142 (2002).

57. Olson, M. F. & Sahai, E. The actin cytoskeleton in cancer cell motility. Clinical and Experimental Metastasis vol. 26, 273–287 (2009).

58. Hodge, R. G. & Ridley, A. J. Regulating Rho GTPases and their regulators. Nature Reviews Molecular Cell Biology vol. 17, 496–510 (2016).

59. Burridge, K., Monaghan-Benson, E. & Graham, D. M. Mechanotransduction: From the cell surface to the nucleus via RhoA. Philosophical Transactions of the Royal Society B: Biological Sciences vol. 374 (2019).

60. Lessey, E. C., Guilluy, C. & Burridge, K. From mechanical force to RhoA activation. Biochemistry 51, 7420–7432 (2012).

61. Jiricny, J. The multifaceted mismatch-repair system. Nature Reviews Molecular Cell Biology vol. 7, 335–346 (2006).

62. Germano, G., Amirouchene-Angelozzi, N., Rospo, G. & Bardelli, A. The Clinical Impact of the Genomic Landscape of Mismatch Repair–Deficient Cancers. Cancer Discov. 8, 1518–1528 (2018).

63. Hanniford, D., et al. Epigenetic Silencing of CDR1as Drives IGF2BP3-Mediated Melanoma Invasion and Metastasis. Cancer Cell 37, 55–70.e15 (2020).

64. Shang, Q., et al. FMRP ligand circZNF609 destabilizes RAC1 mRNA to reduce metastasis in acral melanoma and cutaneous melanoma. J. Exp. Clin. Cancer Res. 41, 170 (2022).

65. Chen, J., et al. Circ-GLI1 promotes metastasis in melanoma through interacting with p70S6K2 to activate Hedgehog/GLI1 and Wnt/β-catenin pathways and upregulate Cyr61. Cell Death Dis. 11, 596 (2020).

66. Mecozzi, N., Vera, O. & Karreth, F. A. Squaring the circle: circRNAs in melanoma. Oncogene vol. 40, 5559–5566 (2021).

67. Shirasaki, R. & Pfaff, S. L. Transcriptional Codes and the Control of Neuronal Identity. Annu. Rev. Neurosci. 25, 251–281 (2002).

68. Matthews, J. M. & Sunde, M. Zinc Fingers-Folds for Many Occasions. IUBMB Life 54, 351–355 (2002).

69. Feuerstein, R., Wang, X., Song, D., Cooke, N. E. & Liebhaber, S. A. The LIM/Double Zinc-Finger Motif Functions as a Protein Dimerization Domain. Proc. Nati. Acad. Sci. USA vol. 91 (1994).

70. Perez-Alvaradol, G. C., et al. Structure of the Carboxy-Terminal LIM Domain from the Cysteine Rich Protein CRP. Nat Struct Biol. 1(6):388–98 (1994).

71. Kadrmas, J. L. & Beckerle, M. C. The LIM domain: From the cytoskeleton to the nucleus. Nature Reviews Molecular Cell Biology vol. 5, 920–931 (2004).

72. Dawid, I. B. & Gov, I. LIM Protein Interactions: Drosophila Enters the Stage. vol. 14 (1998).

73. Herblot, S., Steff, A.-M., Hugo, P., Aplan, P. D. & Hoang, T. SCL and LMO1 Alter Thymocyte Differentiation: Inhibition of E2A-HEB Function and Pre-Tα Chain Expression. Nat Immunol. 1(2):138–44 (2000).

74. Guerrero-Santoro, J., Yang, L., Stallcup, M. R. & DeFranco, D. B. Distinct LIM domains of Hic-5/ARA55 are required for nuclear matrix targeting and glucocorticoid receptor binding and coactivation. J. Cell. Biochem. 92, 810–819 (2004).

75. Drori, S., et al. Hic-5 regulates an epithelial program mediated by PPARγ. Genes Dev. 19, 362–375 (2005).

76. Heitzer, M. D. & DeFranco, D. B. Mechanism of Action of Hic-5/Androgen Receptor Activator 55, a LIM Domain-Containing Nuclear Receptor Coactivator. Mol. Endocrinol. 20, 56–64 (2006).

77. Roof, D. J., Hayes, A., Adamian, M., Chishti, A. H. & Li, T. Molecular Characterization of AbLIM, a Novel Actin-Binding and Double Zinc Finger Protein. The Journal of Cell Biology vol. 138 (1997).

78. Barrientos, T., et al. Two novel members of the ABLIM protein family, ABLIM-2 and-3, associate with STARS and directly bind F-actin. Journal of Biological Chemistry 282, 8393–8403 (2007).

79. Yang, Y., et al. Circular RNA circFCHO2(hsa_circ_0002490) promotes the proliferation of melanoma by directly binding to DND1. Cell Biol. Toxicol. 40, (2024).

80. Te Velthuis, A. J. W. & Bagowski, C. P. PDZ and LIM domain-encoding genes: Molecular interactions and their role in development. The Scientific World Journal vol. 7, 1470–1492 (2007).

81. Zheng, M., Cheng, H., Banerjee, I. & Chen, J. ALP/Enigma PDZ-LIM domain proteins in the heart. Journal of Molecular Cell Biology vol. 2, 96–102 (2010).

82. Beckerle, M. C. Zyxin: Zinc fingers at sites of cell adhesion. BioEssays 19, 949–957 (1997).

